# Deciphering the Multiscale Morphology of Somatic Oncogenic Alterations in Hepatocellular Carcinoma

**DOI:** 10.64898/2026.07.15.738764

**Authors:** Qionghua Shen, Tai Ngo, Hanieh Mazloom-Farsibaf, Kelly Wong, Lin Li, Kushal Bhatt, Felix Y Zhou, Bo-Jui Chang, Xiaoding Wang, Zhiguo Shang, John Haug, Hazel M. Borges, Reto Fiolka, Hao Zhu, Kevin M. Dean

## Abstract

Somatic oncogenic mutations are typically defined by their molecular alterations, yet how they reorganize cellular architecture within intact tissues remain largely unknown. Here, we demonstrate that distinct oncogenic drivers produce unique multiscale architectural phenotypes that can be quantitatively resolved in intact liver tissue. Using iterative expansion microscopy, multiscale light-sheet imaging, and three-dimensional morphometric analysis, we systematically mapped structural remodeling from single-cell morphology to mitochondrial architecture in mosaic mouse models of hepatocellular oncogene activation. NRAS and CTNNB1 induced fundamentally different morphological programs. NRAS activation drove extensive remodeling of cell morphology, membrane curvature, and surface irregularity, whereas CTNNB1 activation largely preserved global cell morphology while selectively altering mitochondrial organization and shape. We first established a zonation-aware reference state for interpreting oncogene-associated organelle remodeling by resolving mitochondrial differences between periportal and pericentral hepatocytes. Within this framework, CTNNB1 activation shifted mitochondrial features toward a pericentral-like state, consistent with the role of Wnt/β-catenin signaling in hepatic zonation and metabolic identity. Furthermore, integrating cellular morphology, membrane geometry, and mitochondrial architecture improved discrimination of oncogenic states beyond any individual structural feature, demonstrating that mutation-specific phenotypes arise through coordinated remodeling across multiple biological scales. Together, these findings establish multiscale structural phenotyping as a framework for linking oncogenic genotype to three-dimensional cellular organization and reveal that distinct oncogenic drivers remodel different architectural compartments during liver oncogene activation.

## Introduction

Compartmentalization is a fundamental feature of eukaryotic cellular organization. Organelles, including the nucleus, mitochondria, endoplasmic reticulum (ER), and Golgi apparatus, create and maintain specialized intracellular environments that support essential cellular functions. Therefore, those organelles are not merely passive cellular components but active regulators of cell fate and function. In cancer, oncogenic remodeling of cellular and subcellular morphology is a hallmark of transformation. For example, mitochondrial fragmentation, membrane deformation, and cytoskeletal reorganization have all been implicated in metabolic adaptation, stress signaling, and invasive behavior during tumor progression^1–4^. As a result, morphology serves as a structural manifestation of the underlying molecular state of a cell. Understanding how oncogenic mutations reshape cellular and subcellular architecture therefore provides an opportunity to connect molecular alterations with their morphological consequences across intact tissue.

Hepatocellular carcinoma (HCC) provides an especially compelling system in which to investigate these relationships. Recent advances in cancer genomics have demonstrated that HCC exhibits substantial molecular heterogeneity^5^. Recurrent alterations in *CTNNB1, TP53*, and RAS/MAPK-associated pathways define biologically distinct HCC subclasses with different metabolic states, microenvironment interactions, and therapeutic responses^6–8^. Importantly, as one of the most organelle rich cells in the body, hepatocytes have highly specialized ER and mitochondrial networks that support the liver’s diverse biosynthetic and metabolic functions.

Because cellular function in hepatocytes is tightly coupled to subcellular organization, perturbations of molecular programs are often accompanied by measurable changes in organelle architecture and cellular morphology. This unique relationship makes the liver an attractive model for investigating how oncogenic mutations are translated into structural phenotypes. However, despite extensive genomic characterization of HCC, the structural consequences of specific oncogenic mutations within intact tissue remain poorly understood. In particular, how specific oncogenic programs remodel organelle architecture, cellular morphology, and tissue organization within intact liver tissue has not been systematically explored. This challenge is further complicated by the intrinsically heterogeneous spatial organization of the liver. In the liver, hepatocytes are organized along periportal-to-pericentral gradients that define distinct metabolic and signaling states in a process known as liver zonation. These spatially patterned states shape both basal organelle organization and the cellular response to oncogenic stress^9^. Consequently, organelle morphology in liver tissue reflects the combined influence of oncogenic signaling and local tissue context. For example, mitochondrial organization is tightly linked to metabolic specialization across liver zonation yet is also profoundly remodeled during HCC development^10–13^.

Determining mutation-associated changes in subcellular organization necessitates the development of imaging approaches that preserve three-dimensional (3D) tissue architecture, and that have sufficient spatial resolution to resolve both intracellular and organelle-level remodeling. However, nanoscale resolution imaging of intact liver tissue is technically challenging because of dense cellular packing, strong light scattering, and intrinsic pigmentation. These properties rapidly degrade imaging depth and contrast. Expansion microscopy (ExM) overcomes this by physically enlarging biological specimens while reducing refractive index heterogeneity, thus improving optical transparency in previously opaque tissues^14,15^. In addition, the physical separation between subcellular structures is increased, enabling conventional diffraction-limited microscopes to resolve nanoscale features. However, expansion also dramatically increases specimen volume, often enlarging tissues to millimeter- or centimeter-scale dimensions. This creates a major imaging challenge: the same process that reveals nanoscale structural information also increases the scale over which imaging must be captured. Light-sheet microscopy is particularly well suited to address this problem because it enables rapid volumetric imaging with optical sectioning, low photobleaching, and high imaging throughput across large 3D samples. Yet linking somatic alterations to cellular and subcellular architecture within intact tissue requires an imaging strategy that bridges nanoscale organelle structure with macroscale tissue organization.

We developed a multiscale imaging framework that integrates iterative expansion microscopy (iExM)^16^ with oblique plane microscopy (OPM) and axially swept light-sheet microscopy (ASLM) to connect tissue-scale organization with nanoscale subcellular architecture within the same intact liver specimen. To address the opposing demands of imaging scale and resolution introduced by ExM, we use OPM for rapid, large-volume mapping of expanded liver tissue and ASLM for targeted high-resolution interrogation of cellular and subcellular structure. OPM enables volumetric imaging across large, expanded specimens, allowing comprehensive mapping of tissue organization and the spatial distribution of oncogene-activated hepatocytes^17^. In parallel, ASLM generates an ultra-thin and axially uniform light sheet that enables isotropic nanoscale imaging of cellular and subcellular structures in expanded tissues at effective biological resolutions approaching tens of nanometers^18,19^. By integrating large-volume tissue mapping with targeted high-resolution nanoscale imaging interrogation, our iExM–OPM–ASLM workflow enables quantitative structural analysis across multiple biological scales, ranging from tissue organization and single-cell morphology to membrane topology curvature and mitochondrial ultrastructure. Using this framework, we investigate how distinct oncogenic drivers remodel liver architecture across spatial scales in intact tissue. We show that multiscale structural phenotyping can distinguish oncogenic states directly from multilevel volumetric morphology, revealing coordinated remodeling of cellular shape, membrane curvature, and mitochondrial architecture associated with distinct oncogenic programs. More broadly, this strategy provides a generalizable platform for linking molecular state to tissue-scale architecture in complex intact biological systems.

## Results

### A dual-mode multiscale imaging platform enables volumetric characterization of liver architecture across tissue, cellular, and subcellular scales

To determine how oncogenic mutations remodel liver tissue, structural changes must be measured across spatial scales while preserving native three-dimensional context. This requires imaging approaches that can map large tissue volumes while also resolving cellular and subcellular architecture within intact specimens. Conventional microscopy platforms are limited in their ability to simultaneously achieve high volumetric throughput, broad spatial coverage, and nanoscale effective resolution in expanded tissues. To overcome these limitations, we developed an integrated multiscale imaging framework that combines iExM with a spatially coordinated dual-mode light-sheet microscopy system (Fig. 1A, Fig S1A). In this workflow, OPM provides rapid volumetric mapping across large, expanded liver specimens, whereas ASLM enables targeted high-resolution imaging of cellular and subcellular architecture. Together, these complementary modalities link tissue-scale organization to nanoscale structural features within the same intact liver specimen.

**Figure 1.**
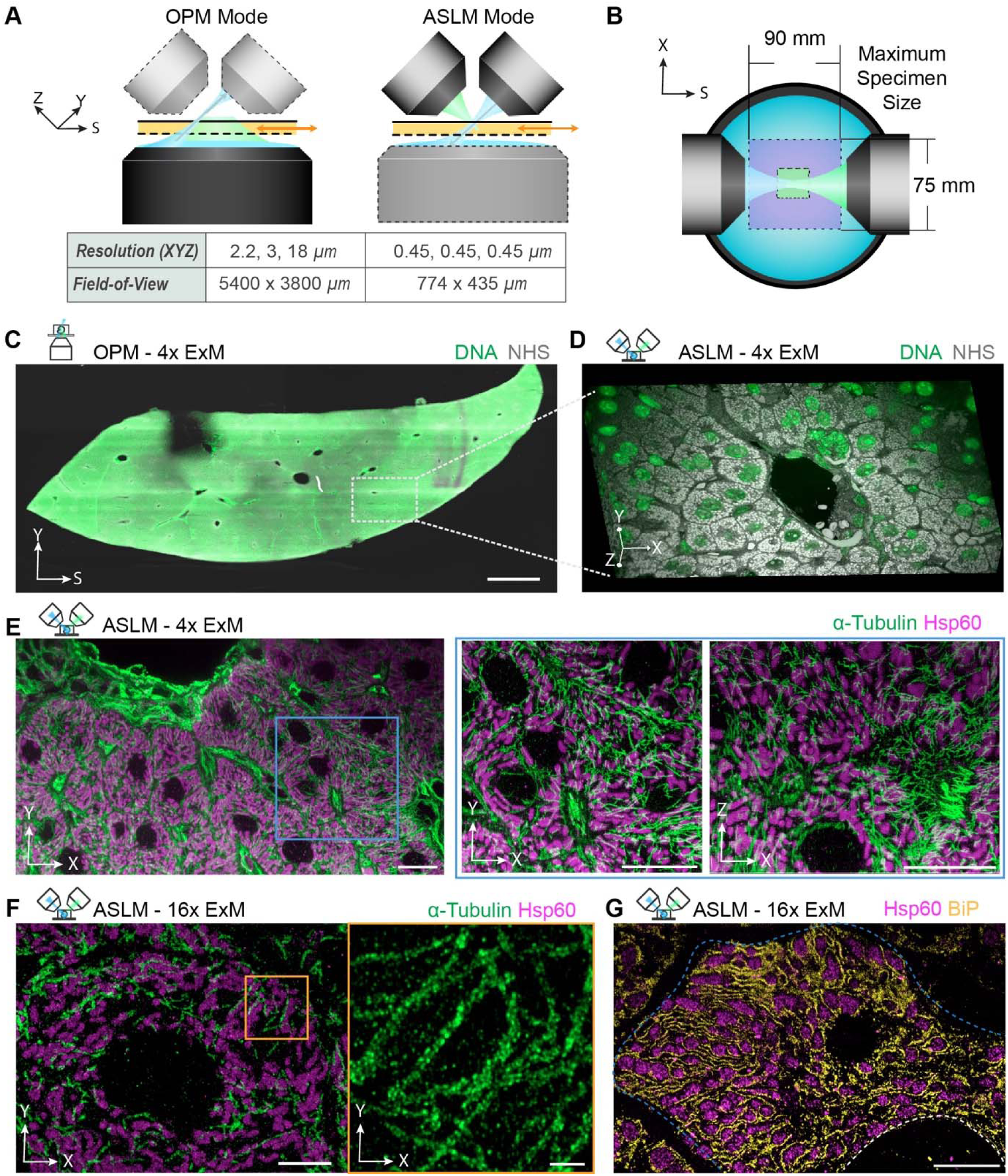
Multiscale imaging platform for rapid volumetric imaging of expanded liver tissue. (A) Three-dimensional schematic of the dual-mode imaging system integrating Oblique Plane Microscopy (OPM, left) and Axially Swept Light-Sheet Microscopy (ASLM, right). The system performance, including spatial resolution and field of view (FOV), is summarized in the table below. (B) Imaging path and sample holder design. The maximum sample size is determined by the holder geometry, the current configuration supports samples up to 90 mm x 75 mm. (C–D) Spatial coordination between OPM and ASLM imaging modalities. An intact liver specimen stained with SYTOX Green (nuclei) and NHS-amine (pan-cellular labeling) was imaged using both systems. (C) Large-scale volumetric imaging of an intact liver specimen acquired with bottom-view OPM using a 4x objective. The imaged volume spans 3.75 mm x 9.8 mm with a depth of 400 µm (post 4x expansion), revealing overall hepatic lobular organization. Scale bar, 4000 µm (post-expansion; equivalent to 1000 µm pre-expansion). (D) Zoom-in three-dimensional rendering of a vascular region from (C), highlighting local tissue architecture. The FOV is 783.36 µm x 440.64 µm x 126 µm (post 4x expansion). (E) ASLM imaging of a 4x expanded liver specimen labeled for α-tubulin (green) and mitochondria (magenta). Right panels show corresponding xy and zx orthogonal views from the region indicated in the left panel (blue box), demonstrating three-dimensional subcellular resolution. Scale bar, 50 µm (post-expansion; equivalent to 12.5 µm pre-expansion). (F) ASLM imaging of a 16x expanded liver specimen labeled for α-tubulin (green) and mitochondria (magenta). Right panels show zoom in regions from the left α-tubulin image, illustrating enhanced effective resolution at higher expansion factors. Left scale bar, 50 µm (post-expansion; equivalent to 3.12 µm pre-expansion); right scale bar, 10 µm (post-expansion; equivalent to 0.625 µm pre-expansion). (G) High-resolution 16× expansion imaging of liver tissue illustrating subcellular interactions between mitochondria (pink) and endoplasmic reticulum (yellow). Dashed blue line indicates the cell boundary and dashed white line indicates the nuclei. Scale bars, 50 μm (post-expansion) and 12.5 μm (pre-expansion).

We first adapted the iExM workflow for thick liver specimens to achieve reproducible, isotropic physical magnification while preserving tissue architecture and subcellular organization. Fixed 100-µm liver sections were processed through two sequential rounds of hydrogel embedding and isotropic swelling, producing linear expansion factors of approximately 4x after the first round and up to 16x after the second round (Fig. S1A–F). This stepwise expansion increased the effective separation between intracellular structures without requiring changes to the optical configuration of the microscope, enabling the same platform to operate across distinct effective resolution regimes. Validation across tissue-scale and nuclear features showed preserved morphology after successive expansion rounds, including clear visualization of subnuclear structures such as nucleoli (red arrows, Fig. S1E). Thus, the modified iExM workflow provides a robust sample-preparation foundation for linking tissue-scale imaging with nanoscale structural interrogation in expanded liver tissue.

To enable multiscale imaging of expanded liver tissues, we customized a dual-mode light-sheet platform that integrates OPM and ASLM (Fig. 1A–B). The system was designed to combine rapid, large-volume tissue mapping with targeted high-resolution imaging, thereby linking tissue-scale spatial organization to nanoscale subcellular morphology within expanded intact tissue specimens. The OPM imaging mode enabled rapid volumetric mapping of expanded liver tissue across millimeter-scale fields of view. In this configuration, the OPM provides a ∼5.4 x 2.0-mm optical field of view, while larger lateral extents can be accessed by sample scanning and were limited primarily by the sample chamber and coverslip dimensions (∼75 × 90 mm). The field-of-view and resolution of this mode is sufficient to resolve hepatic lobular organization, large vascular structures, and the spatial distribution of labeled cells across expanded tissue volumes (Fig. 1A, C). In parallel, the ASLM mode provided isotropic high-resolution volumetric imaging with a raw resolution of ∼0.45 µm, corresponding to an estimated effective biological resolution of ∼100 nm after 4x expansion and ∼30 nm after 16x expansion (Fig. 1A, D–F). This resolution regime enables visualization of cellular, intracellular, and organelle-level architecture within intact expanded tissue specimens. Importantly, the OPM and ASLM imaging modes can be spatially registered, allowing regions identified during large-volume OPM imaging to be revisited by ASLM without losing tissue coordinates. This coordinate-preserving workflow enabled rapid transitions between tissue-scale mapping and targeted subcellular interrogation. As a demonstration, intact expanded liver tissues labeled with SYTOX Green and NHS-ester were first imaged by OPM to reconstruct large-scale tissue organization, revealing preserved hepatic lobular structures and spatial heterogeneity across the tissue volume (Fig. 1C). Regions of interest identified from OPM datasets were then imaged using ASLM, to reveal local cellular organization, and intracellular morphology within the same tissue coordinate system (Fig. 1D).

To further evaluate the subcellular resolving capability of the platform, liver tissues labeled for α-tubulin and Hsp60 (mitochondria) were imaged under both 4x and 16x expansion conditions (Fig. 1E–F). At 4x expansion, ASLM resolved abundant mitochondrial networks and complex, interdigitating microtubules throughout intact hepatocytes (Fig. 1E). These datasets also revealed regional variation in cytoskeletal organization, with hepatocytes and vessel-associated cells exhibiting distinct microtubule patterns within the same tissue region. Increasing the expansion factor to 16x further enhanced effective biological resolution, enabling visualization of fine mitochondrial features and dense subcellular filament organization that were indistinguishable at the lower expansion (Fig. 1F). This increased nanoscale resolution further revealed extensive spatial interactions between mitochondria and endoplasmic reticulum networks within hepatocytes (Fig. 1G). Together, these results demonstrate that the combined iExM–OPM–ASLM workflow enables coordinated structural interrogation across tissue, cellular, and organelle scales, providing a platform for quantifying how liver architecture and intracellular organization are remodeled within intact tissue.

### Multiscale imaging reveals oncogene-specific single-cell morphological remodeling in intact liver tissue

Building on the iExM–OPM–ASLM imaging framework, we next used this integrated platform to characterize oncogene-driven cellular remodeling within intact liver tissue. Mosaic oncogene activation was generated by hydrodynamic delivery of transposon constructs encoding either NRAS or CTNNB1, producing spatially distributed oncogene-activated hepatocytes throughout the liver (Fig. 2A). Volumetric OPM imaging enabled visualization of these hepatocytes across intact expanded tissue volumes (Fig. 2A, B). Consistent with analysis of zonal position, cells overexpressing oncogenes were broadly distributed across hepatic lobules rather than confined to a specific periportal or pericentral region (Fig. S4A). This mosaic distribution provided a spatially heterogeneous tissue context for evaluating oncogene-specific cellular remodeling.

**Figure 2.**
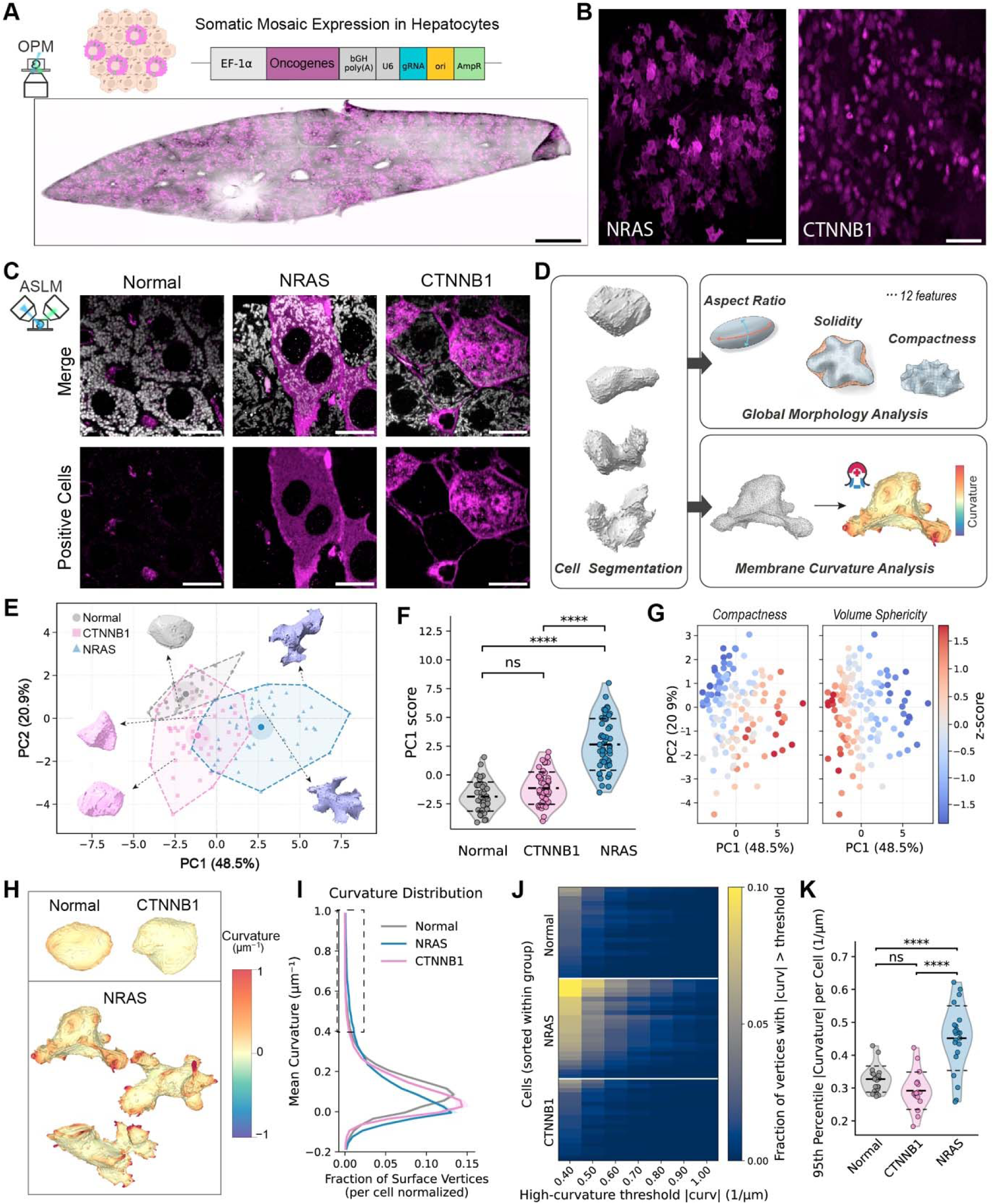
Single-cell morphological remodeling following oncogenic activation in liver tissue. (A) Whole specimen view of liver tissue imaged by bottom-view Oblique Plane Microscopy (OPM), illustrating the spatial distribution of oncogene activated cells. The top panel shows the plasmid design and mosaic activation strategy used to induce sparse oncogenic expression. Scale bar, 4000 µm (post-expansion; equivalent to 1000 µm pre-expansion). (B) Zoomed-in OPM views of liver regions containing distinct oncogene activated populations (magenta). These images highlight spatial heterogeneity and local tissue organization following activation. Scale bar, 500 µm (post-expansion; equivalent to 125 µm pre-expansion). (C) High-resolution imaging of individual hepatocytes from normal, NRAS, and CTNNB1 activated tissues acquired using Axially Swept Light-Sheet Microscopy (ASLM). Cell structures are shown in magenta, revealing detailed morphological differences at the single-cell level. Scale bar, 50 µm (post-expansion; corresponding pre-expansion 12.5 µm accordingly). (D) Workflow for single-cell morphological analysis. The left panel shows representative three-dimensional renderings of segmented individual cells. The right panels illustrate quantitative analysis pipelines, including global geometric feature extraction (top) and membrane curvature analysis (bottom). (E) Principal component analysis (PCA) of single-cell morphological features across three groups: Normal (gray, n = 42), CTNNB1 (pink, n = 50), and NRAS (blue, n = 49). Each point represents an individual cell. Insets show representative cell morphologies for each group. Large double-circle markers indicate group centroids, and polygons outline the distribution of each population in feature space. (F) Statistical comparison of PC1 scores across the three groups, highlighting shifts in global morphology following oncogenic activation (p**** < 0.0001, ns = 0.11). (G) Distribution of selected global morphological features, including compactness (left) and volume sphericity (right), across Normal, CTNNB1, and NRAS groups. (H) Representative mesh maps of membrane curvature for individual cells from each group, illustrating differences in surface topology. (I) Distribution of membrane curvature values for each group, normalized at the single-cell level. The dashed region highlights the high-curvature tail of the distribution. (J) Heatmap showing the frequency of high-curvature regions across cells for each condition, corresponding to the tail region defined in (I). (K) Quantitative comparison of high-curvature metrics (95% tail area) between groups, demonstrating mutation-specific differences in membrane surface irregularity (p**** < 0.0001, ns = 0.2862).

High-resolution ASLM imaging revealed distinct three-dimensional morphologies among normal, NRAS-activated, and CTNNB1-activated hepatocytes. NRAS-activated hepatocytes exhibited pronounced morphological remodeling, with irregular cell shapes, elongated protrusions, and increased surface roughness. In contrast, CTNNB1-activated hepatocytes retained comparatively smooth and compact morphologies that more closely resembled normal hepatocytes (Fig. 2C). To quantify these phenotypes, individual hepatocytes were segmented from volumetric datasets and analyzed using a multiscale geometric analysis pipeline combining 13 quantitative global morphology features (including sphericity, solidity, aspect ratio, compactness, roughness, surface area, and volume, see Supplementary Table 1) and membrane curvature measurement (Fig. 2D).

Principal component analysis (PCA) of global morphology features separated normal, NRAS-activated, and CTNNB1-activated hepatocytes (Fig. 2E). Normal and CTNNB1-activated cells largely overlapped, occupying a region associated with smooth, rounded morphologies, whereas NRAS-activated cells formed a distinct cluster characterized by increased shape irregularity and protrusive shapes (Fig. 2E). The first principal component (PC1), accounting for 48% of the total variance, represented the dominant axis of morphological variation, distinguishing spherical from angular cell morphologies (Fig. 2F). Examination of feature loadings revealed that higher PC1 scores were associated with increased surface irregularity, reduced sphericity, and more protrusive cell shapes, consistent with the distribution of NRAS-activated hepatocytes, whereas lower PC1 scores reflected the smoother, more blob-like morphologies observed in normal and CTNNB1-activated cells (Fig. 2G). Feature-level comparisons further supported these mutation-specific structural signatures, identifying NRAS activation as a major driver of global morphological remodeling, while CTNNB1 activation produced comparatively subtle changes that remained largely similar to the normal hepatocyte population. Together, these findings demonstrate that distinct oncogenic drivers induce reproducible and mutation-specific alterations in hepatocyte morphology.

Analysis of cell membrane curvature revealed NRAS-activated hepatocytes contained extensive high-curvature membrane regions distributed across the cell surface, consistent with their protrusive and irregular morphologies (Fig. 2H). In contrast, CTNNB1-activated hepatocytes exhibited more uniform curvature distributions, similar to normal hepatocytes. Population-level curvature analysis demonstrated enrichment of high-curvature membrane domains in NRAS-activated cells (Fig. 2I, J), a pattern also evident in representative single-cell curvature histograms (Fig. S4C). Quantification of the 95th percentile curvature confirmed that NRAS activation significantly increased membrane curvature compared with both normal and CTNNB1-activated hepatocytes (Fig. 2K). Together, these findings show that distinct oncogenic drivers generate quantitatively separable three-dimensional cell morphologies and membrane curvature states within intact liver tissue.

### Multiscale imaging establishes a zonation-aware mitochondrial reference state for evaluating oncogene-associated remodeling

Having established that oncogenic drivers induce divergent single-cell morphologies, we next asked whether these morphological differences extend to the subcellular level, for example mitochondrial architecture. Mitochondria are central to hepatocyte metabolism, and their shapes and organization are tightly coupled with metabolic state and liver zonation. We therefore first tested whether the iExM–OPM–ASLM workflow could resolve known differences in mitochondrial organization between periportal and pericentral hepatocytes, establishing a physiological benchmark for subsequent analysis of oncogene-associated remodeling (Fig. 3A).

**Figure 3.**
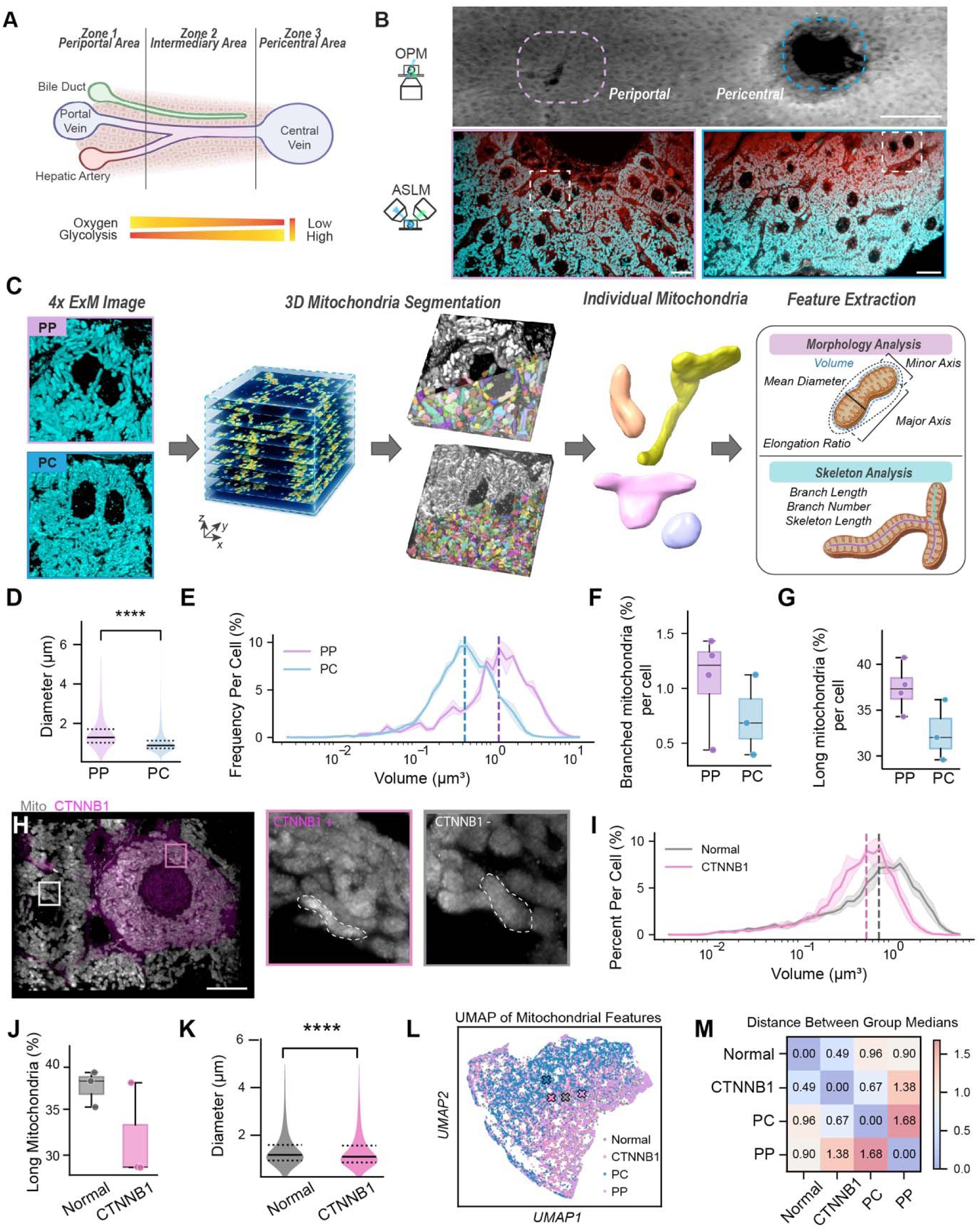
Mitochondrial morphological quantification across liver zonation and oncogenic activation. (A) Schematic of physiological liver zonation, illustrating spatial gradients of oxygen and metabolic activity (glycolysis). (B) Immunofluorescence (IF) imaging reveals periportal (PP) and pericentral (PC) regions; top, whole-tissue view acquired by OPM; bottom, high-resolution ASLM images showing tissue to subcellular (mitochondrial) organization (left, PP; right, PC). Scale bars: 500 μm (top), 50 μm (bottom) post-expansion, corresponding to 125 μm and 12.5 μm pre-expansion. (C) Workflow for mitochondrial morphological quantification, including single-cell extraction, 3D segmentation, and feature analysis based on global morphology (top) and skeletonization (bottom). (D) Quantification of mean mitochondrial diameter in PP (n = 2,197) and PC (n = 4,895) regions (p **** < 0.0001). (E–G) Quantification of mitochondrial volume distribution (cell-normalized), branching percentage, and proportion of long mitochondria (top third, length > 1.2 μm) across PP and PC regions. (H) IF images show altered mitochondrial organization in CTNNB1-activated cells (pink) compared to non-activated normal cells (gray). Dashed line indicates the boundary of a single mitochondria. Scale bar: 50 μm post-expansion equivalent to 12.5 μm pre-expansion. (I–K) Quantification of mitochondrial volume distribution (cell-normalized), long mitochondria proportion (length > 2.3 μm), and mean diameter in Normal (n = 6,715) and CTNNB1⁺ (n = 6,269) cells (p **** < 0.0001). (L) UMAP embedding of mitochondrial features across four groups (Normal, CTNNB1⁺, PC, PP), with crosses indicating group medians. (M) Pairwise distance heatmap of group medians across all mitochondrial features. For all the quantification, n ≥ 3 cells each group.

Combined OPM and ASLM imaging revealed pronounced spatial differences in mitochondrial organization across intact liver lobules (Fig. 3B). OPM enabled identification of periportal and pericentral regions within expanded liver tissue, while ASLM resolved mitochondrial morphology within these regions at subcellular resolution following 4x expansion (Fig. S5A–C). Periportal hepatocytes contained larger and more elongated mitochondrial structures, whereas pericentral hepatocytes contained smaller and less elongated mitochondrial structures. To quantify these differences, we developed an analysis workflow integrating three-dimensional segmentation, single-object extraction, and their volume-based (including mean diameter, volume, major and minor axis length, elongation ratio) and skeleton-based (branch number, total skeleton length) morphometrics (Fig. 3C).

Our quantitative analysis confirmed zonation-dependent differences in mitochondrial architecture. Periportal hepatocytes contained mitochondrial structures with larger mean diameters than pericentral hepatocytes (median, 1.29 µm versus 0.88 µm; Fig. 3D). Mitochondrial volume distributions were similarly shifted toward larger segmented objects in periportal regions (median, 1.15 µm³ versus 0.42 µm³; Fig. 3E). Skeleton-based analysis further showed that periportal hepatocytes had higher occurrence of mitochondrial branching longer mitochondria (Fig. 3F, G). Together, these measurements indicate that periportal hepatocytes contain larger, connected, and more highly branched mitochondrial structures than pericentral hepatocytes. These results agree with previous reports of metabolic and ultrastructural zonation of hepatic mitochondria and demonstrate that our platform can resolve physiologically relevant subcellular heterogeneity within intact liver tissue^12,13^. Importantly, our analysis establishes a zonation-aware mitochondrial reference state for evaluating oncogene-associated remodeling.

### CTNNB1 activation shifts mitochondrial architecture toward a pericentral-like state

Having established that our workflow captures baseline mitochondrial zonation, we next asked whether oncogenic signaling remodels mitochondrial architecture relative to this physiological reference state. We focused primarily on CTNNB1 activation because Wnt/β-catenin signaling is a central regulator of pericentral hepatocyte identity and metabolic zonation, providing a direct link between an oncogenic driver and a spatially patterned mitochondrial state. ASLM imaging revealed that CTNNB1-activated hepatocytes contained smaller and less elongated mitochondrial structures relative to neighboring normal hepatocytes within the same zonal region (Fig. 3H). Quantitative analysis showed that CTNNB1 activation shifted mitochondrial volume distributions toward smaller objects and reduced the fraction of long mitochondria (median mitochondrial volume, 0.513 µm³; median mean diameter, 1.09 µm; Fig. 3I–K). All comparisons were performed within matched liver zonation regions to minimize confounding from baseline periportal–pericentral mitochondrial heterogeneity. Because analyses were performed at the earliest stages of oncogenic activation, prior to detectable changes in tissue-scale liver zonation, the observed differences primarily reflect oncogene-associated structural remodeling rather than zonal reprogramming. To investigate whether this mitochondrial phenotype reflected a broader consequence of oncogenic activation or a driver-specific effect, we performed the same analysis on NRAS-activated hepatocytes. In contrast to CTNNB1 activation, NRAS activation did not produce significant changes in mitochondrial volume, branching, or length relative to neighboring normal hepatocytes from matched zonal regions (Fig. S6A, C, D). Thus, despite the pronounced cell-shape remodeling induced by NRAS activation, mitochondrial architectural remodeling was more strongly associated with CTNNB1 activation in the features measured here.

To determine how CTNNB1-associated mitochondrial remodeling relates to physiological zonation, we compared mitochondrial feature states across normal, CTNNB1-activated, periportal, and pericentral hepatocytes (Fig. 3L). UMAP visualization showed that CTNNB1-associated mitochondrial architecture shifted toward the pericentral state, consistent with a smaller and less elongated mitochondrial phenotype. Pairwise distances between group medians computed across all mitochondrial features supported this relationship, with CTNNB1-activated mitochondria positioned closer to pericentral than periportal mitochondrial states (Fig. 3M). This finding is consistent with the established role of endogenous Wnt/β-catenin signaling in maintaining pericentral hepatocyte identity and metabolic zonation within the normal liver lobule^20,21^. Together, these results indicate that CTNNB1 activation biases mitochondrial morphology toward a pericentral-like structural state, whereas NRAS-driven transformation produces comparatively limited remodeling of mitochondrial morphology. More broadly, this analysis links oncogenic signaling to spatially patterned subcellular organization within intact liver tissue.

### Integration of multiscale structural features enables prediction of oncogenic states

After establishing that NRAS and CTNNB1 activation induce distinct structural phenotypes at cellular and subcellular scales, we next asked whether these architectural features could be computationally integrated to infer oncogenic state directly from intact tissue morphology. We therefore concatenated quantitative descriptors of global cell morphology, membrane curvature, and mitochondrial shape into a unified multiscale analysis framework (Fig. 4A, B). These feature classes captured complementary aspects of oncogene-associated remodeling: NRAS-activated hepatocytes exhibited increased membrane curvature and shape irregularity, whereas CTNNB1-activated hepatocytes showed comparatively modest global shape disruption but distinct mitochondrial architectural remodeling. Feature-wise z-score normalization was then used to visualize relative enrichment of each structural feature across normal, NRAS-activated, and CTNNB1-activated hepatocytes (Fig. 4B).

**Figure 4.**
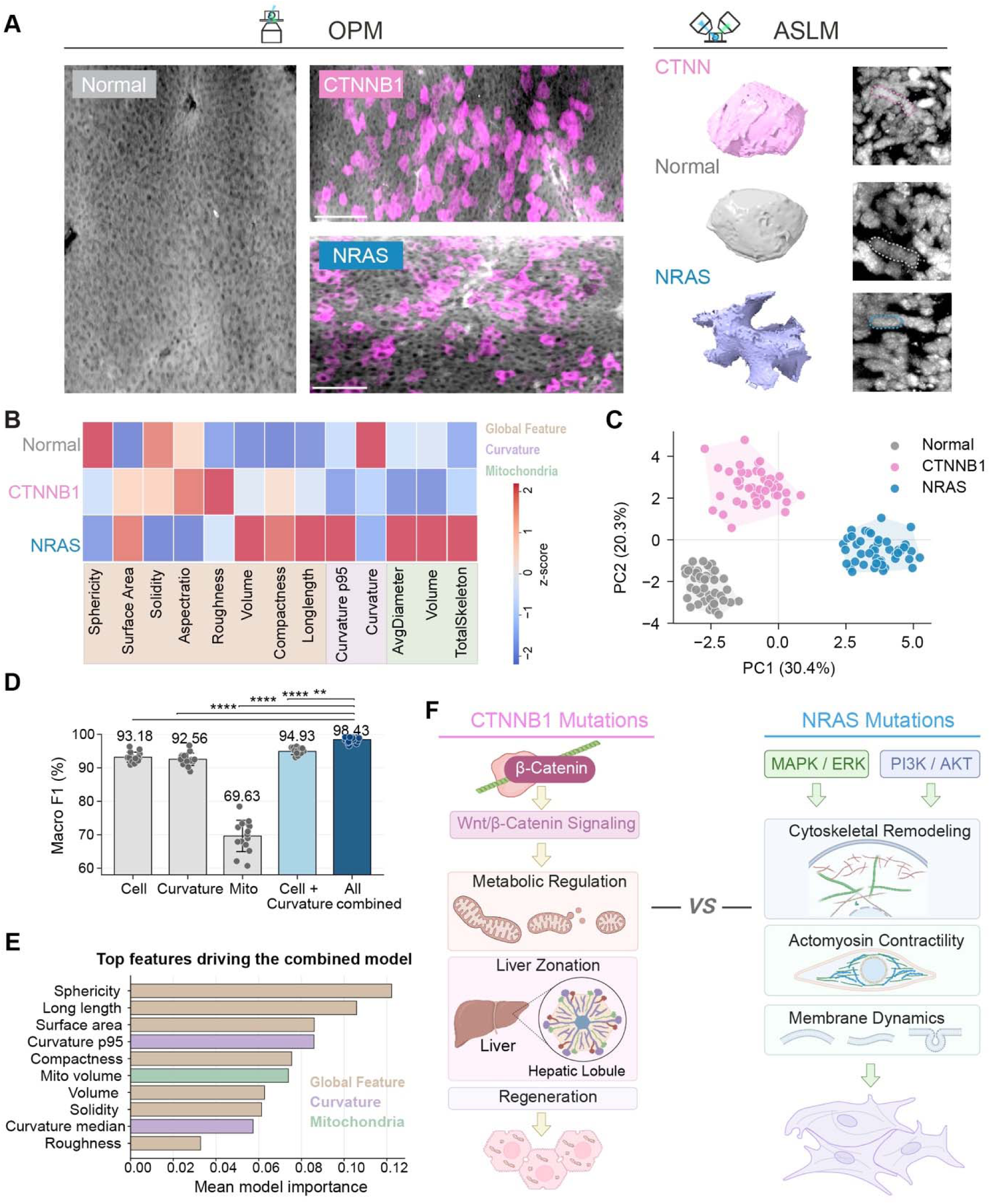
Multiscale integration of cellular and subcellular features enables discrimination of oncogenic states. (A) Multiscale IF imaging showing tissue wide organization, single cell morphology, and subcellular mitochondrial structure in Normal (gray), NRAS-activated (blue), and CTNNB1-activated (pink) hepatocytes. Scale bar, 500 μm (post-expansion) and 125 μm (pre-expansion). (B) Heatmap of z-scored group means for global cell morphology, membrane curvature, and mitochondrial features across the three groups. (C) Principal component analysis (PCA) of integrated pseudo-bulk feature vectors. Each point represents a pseudo-bulk sample generated by random aggregation of cell morphology, curvature, and mitochondrial measurements within each group (n = 50 per group). Polygons indicate group distributions. (D) Classification performance (macro F1 score) across feature sets (cell morphology, curvature, mitochondria, and combinations), evaluated using GroupKFold cross-validation with repeated sampling. Dots represent individual repeats and bars indicate mean ± s.d. p-values for comparisons with the combined model are: Cell, p ****< 0.0001; Curvature, p ****< 0.0001; Mitochondria p **** < 0.0001; Cell + curvature, p ** < 0.01. (E) Ranked feature importance from the combined model, showing contributions from global morphology, curvature, and mitochondrial features. (F) Schematic summarizing differential contributions of CTNNB1 and NRAS activation to hepatic tissue remodeling.

Because structural measurements were acquired across different biological scales, we generated genotype-matched pseudo-bulk profiles to integrate cellular, membrane-curvature, and mitochondrial features into a common feature space. Each pseudo-bulk profile combined randomly sampled measurements from the same genotype across the three feature sets, summarized by modality-level statistics that captured both average structural state and within-group variability (Fig. S6H). These aggregated features were concatenated into unified multiscale vectors and standardized before dimensionality reduction. Principal component analysis of the integrated pseudo-bulk profiles revealed clear separation among normal, CTNNB1-activated, and NRAS-activated populations (Fig. 4C). Each point in this PCA representation therefore corresponds to one integrated multiscale structural profile of a spatially patterned pseudo-cell. These results show that coordinated analysis of cellular morphology, membrane curvature, and mitochondrial architecture captures genotype-specific structural states that are not apparent from any single scale alone.

We next tested whether these multiscale structural profiles could predict oncogenic state. Supervised classifiers were trained using individual feature sets or the concatenated multiscale feature set and evaluated by repeated grouped cross-validation (Fig. 4D). Global cell-morphology features alone enabled discrimination among normal, NRAS-activated, and CTNNB1-activated hepatocytes (F1 score: 93.18%), while membrane-curvature measurements added complementary discriminatory information (F1 score: 94.93%). Mitochondrial features were less predictive when used alone (F1 score: 69.63%) but improved performance when combined with cellular morphology and curvature measurements (F1 score: 98.43%). This indicates that cellular, membrane, and mitochondrial features encode complementary, potentially identifying, information about oncogenic state. Feature-importance analysis further identified sphericity, long-axis length, surface area, curvature metrics, and mitochondrial volume as major contributors to classification performance (Fig. 4E). These discriminative features aligned with the driver-specific phenotypes observed above: CTNNB1 activation produced comparatively modest global cell-shape disruption but pronounced mitochondrial remodeling, whereas NRAS activation was associated with increased membrane curvature, shape irregularity, and protrusive cell morphology (Fig. 4F). Thus, integrated volumetric structural phenotyping holistically describes mutation-specific architectural states in intact tissue linking oncogenic signaling programs to coordinated remodeling across cellular and subcellular scales.

## Discussion

A central finding of this study is that structural organization across multiple biological scales represents an important layer of phenotype through which oncogenic programs are manifested in intact tissues. Although not all oncogenic alterations are expected to produce readily detectable architectural changes, the distinct oncogenic drivers examined here were associated with separable patterns of structural remodeling across cellular and subcellular compartments. CTNNB1 activation was primarily associated with mitochondrial architectural remodeling, consistent with the established role of Wnt/β-catenin signaling in hepatic zonation and metabolic regulation, whereas NRAS activation predominantly altered cellular morphology, producing increased membrane curvature, irregularity, and protrusive cell shapes^8,22,23^. Importantly, integrating that structural information across cellular and subcellular scales provided greater predictive power than any individual feature class alone. This observation indicates that oncogenic state is not encoded within a single morphological hallmark but is instead distributed across interconnected architectural features spanning multiple biological scales. More broadly, these findings support the concept that tissue architecture contains biologically meaningful information about molecular and genetic state and that multiscale structural phenotyping can provide a quantitative framework for linking genotype to phenotype in intact tissues. Unlike conventional computational pathology approaches, which predominantly rely on two-dimensional histological features^24–26^, the framework presented here directly quantifies three-dimensional tissue organization, cellular morphology, membrane topology, and organelle architecture. As a result, predictive structural signatures can be related to specific biological processes, including cytoskeletal remodeling, membrane dynamics, mitochondrial organization, and zonated metabolic identity. In this sense, multiscale structural phenotyping complements spatial genomics and proteomics by providing an architectural layer of information that connects molecular programs to the physical organization of cells and organelles within native tissue environments.

These biological insights were enabled by the integrated imaging and analysis framework developed in this study. A longstanding challenge in cancer biology is to connect molecular perturbations with their structural consequences while preserving tissue context. Dissociation-based approaches provide rich molecular information but disrupt native cellular architecture, whereas conventional spatial imaging methods often lack the spatial resolution and volumetric coverage required to quantify remodeling across cellular and subcellular scales^27^. Although electron microscopy remains the benchmark for ultrastructural imaging, its limited throughput, restricted field of view, and demanding sample preparation make it difficult to apply across large tissue volumes and biological replicates. By combining iterative expansion microscopy with complementary light-sheet imaging modalities and quantitative morphometric analysis, our framework bridges tissue-scale spatial context with nanoscale structural measurements within the same specimen. This enables systematic characterization of how oncogenic programs are translated into the three-dimensional organization of cells and organelles within intact tissues.

Several limitations should be considered. First, this study focused on two oncogenic drivers and an early stage of oncogenic activation. Whether the structural signatures identified here generalize across additional mutations, combinations of oncogenic events, or later stages of tumor progression remains unclear. Second, although mitochondrial morphology provided a biologically informative subcellular readout, integrating this framework with functional metabolic imaging, biosensors, or spatial metabolomics would more directly connect structural remodeling with cellular function. Third, the multiscale integration relied on pseudo-cell profiles assembled from independently acquired measurements across biological scales. Future studies achieving fully registered tissue-, cell-, and organelle-level measurements from the same cells will provide a more direct view of how oncogenic signaling propagates across biological scales.

Beyond these limitations, an important conceptual distinction is that our findings identify the structural consequences of oncogenic signaling rather than the structural determinants of tumorigenesis. The architectural signatures described here do not establish whether specific morphological features directly contribute to transformation, invasion, or disease progression. Instead, they demonstrate that oncogenic activation generates reproducible and measurable structural states in native tissue context. This distinction is critical because it positions structural phenotyping as a framework for mapping genotype-to-architecture relationships, while future perturbation-based studies will be required to determine which structural features are functionally required for tumor initiation, progression, or therapeutic response.

Looking forward, combining multiscale structural phenotyping with spatial transcriptomics, spatial proteomics, and functional imaging offers an opportunity to construct comprehensive maps linking molecular state, cellular architecture, and tissue organization. Expanding the oncogenic driver panel, incorporating additional organelle systems such as the endoplasmic reticulum, lipid droplets, peroxisomes, bile canalicular networks, and ER–mitochondria contact sites, and extending the analysis to longitudinal models and human HCC specimens will help determine how broadly structural states can classify disease progression and predict oncogenic programs. Together, these findings establish multiscale structural phenotyping as a framework for understanding how oncogenic signaling is translated into the three-dimensional organization of cells and organelles within intact tissues and position cellular architecture as a quantitative phenotype for studying cancer initiation, progression, and therapeutic response.

## Methods

### Mouse liver oncogene activation and tissue preparation

All animal experiments were approved by the IACUC at The University of Texas Southwestern Medical Center under protocol 2017-102315-CORE and performed in accordance with institutional guidelines. Somatic mosaic oncogene activation in hepatocytes was achieved using hydrodynamic tail vein delivery (HDT) in wild-type male C57BL/6 mice (JAX Stock No. 000664) at 8 weeks of age. Only male mice were used to minimize variability associated with sex-dependent liver metabolic differences. HDT enables rapid delivery of transposon and transposase plasmids into hepatocytes, resulting in genomic integration and mosaic overexpression of oncogenic drivers in approximately 1–5% of hepatocytes. Two common hepatocellular carcinoma (HCC) oncogenic drivers, CTNNB1 and NRAS, were evaluated in this study. Mice were injected with 1 µg of Sleeping Beauty transposase plasmid (SB100) together with 10 µg of pT3 transposon vectors encoding either CTNNB1^N90^ or NRAS^G12V^ or empty vector as control (Normal).

Four days after HDT injection, livers were harvested and fixed in 10% formalin for 24 h before transferring to PBS for downstream processing. Fixed liver tissues were embedded in 2% agarose and sectioned into 100 µm-thick slices. Heat-induced epitope retrieval was subsequently performed using Tris-EDTA antigen retrieval buffer (Enzo Life Sciences, ENZ-ACC113) at 95 °C for 30 min. Tissue sections were then washed three times in PBS for 30 min each. And blocked for 2 h at room temperature in blocking buffer containing 4% bovine serum albumin (BSA; Equitech-Bio, BAH65) and 0.2% Triton X-100 (Sigma-Aldrich, X100-100ML) in PBS. Primary antibodies were diluted in blocking buffer and applied to tissue sections for 24 h at 4 °C with gentle agitation. The following primary antibodies were used: anti-Hsp60 (Abcam, ab59457, RRID: AB_2121285, 1:100), anti-β-Catenin (Cell Signaling Technology, D10A8 XP®, 8480, RRID: AB_11127855, 1:100), anti-β-Catenin (Cell Signaling Technology, 37447, RRID: AB_3246427, 1:100), anti-NRAS (Proteintech, 10724-1-AP, RRID: AB_2154209, 1:100), anti-Tubulin (Sigma-Aldrich, T9026, RRID: AB_477593, 1:100), and anti-GRP78 BiP (Abam, ab21685, RRID: AB_2119834, 1:100). Sections were washed three times in PBST for 1 h each after primary incubation. Secondary antibodies, including goat anti-rabbit IgG Alexa Fluor™ 568 (Thermo Fisher Scientific, A-11036) and chicken anti-mouse IgG Alexa Fluor™ 488 (Invitrogen, A-21200), were diluted 1:100 in blocking buffer and incubated with samples overnight at 4 °C. After secondary antibody incubation, tissues were washed three times in PBST for 1 h each. For iterative expansion microscopy experiments, an additional round of secondary antibody labeling was performed to further amplify fluorescence signal intensity before downstream expansion and imaging. Staining specificity and reliability were validated using blank controls as negative controls and double-mutant tissue samples as double-positive controls (Fig. S4B). Positive cells were identified based on transgene fluorescence, with a threshold of 500 arbitrary intensity units (a.u.) above the local background. Cell identification was therefore not performed in a blinded manner.

### Expansion Microscopy

Tissue sections were incubated overnight at room temperature in PBS containing 0.1 mg/mL Acryloyl-X, SE (Thermo Fisher Scientific) to covalently anchor biomolecules to the hydrogel matrix. After two 15 min PBS washes, samples were incubated overnight at 4 °C in monomer solution containing 19% (w/w) sodium acrylate, 10% (w/w) acrylamide, and 0.1% (w/w) N,N′-methylenebisacrylamide. For hydrogel embedding, samples were transferred into custom gelation chambers assembled from microscope slides and silicone spacers (McMaster-Carr, 6459N112) and immersed in freshly prepared polymerization solution supplemented with 0.5% (w/w) ammonium persulfate (APS) and 0.5% (w/w) tetramethylethylenediamine (TEMED). Polymerization was initiated on ice for 30 min and completed at 37 °C for 1.5 h in a humidified chamber. Following polymerization, gels were homogenized in digestion buffer containing 50 mM Tris (pH 8.0), 1 mM EDTA, 0.5% Triton X-100, 1 M NaCl, and 8 U/mL Proteinase K (MilliporeSigma, 39450-01-6) at 37 °C for 6 h. Gels were subsequently washed three times in PBS for 30 min each and labeled with Atto 647 NHS ester (Sigma, 07376-1MG-F; 1:1000 in PBS) for pan-protein visualization and with or without SYTOX™ Green Nucleic Acid Stain (Thermo Fisher Scientific; 1:3000 in PBS) for nuclear labeling. After additional PBS washes (3 x 1 h), samples were expanded into deionized (DI) water with three exchanges of fresh water before imaging, yielding an average linear expansion factor of approximately 4x.

For iterative 16x expansion microscopy, primary expanded gels underwent sequential re-embedding and re-expansion steps following a modified protocol adapted from Deblina et al.^16^. Expanded gels were first equilibrated four times in re-embedding solution containing 13.75% (w/v) acrylamide, 0.038% (w/v) bis-acrylamide, 0.025% (w/v) APS, and 0.025% (w/v) TEMED for 30 min per incubation at room temperature with gentle agitation. Gels were then re-polymerized in nitrogen-flushed chambers at 45 °C for 2 h and washed three times in PBS. For the final expansion step, re-embedded gels were incubated four times on ice in a third gelation solution containing 8.625% (w/v) sodium acrylate, 2.5% (w/v) acrylamide, 0.038% (w/v) bis-acrylamide, 0.025% (w/v) APS, and 0.025% (w/v) TEMED. Polymerization was performed in nitrogen-flushed chambers at 60 °C for 1.5 h, followed by extensive PBS washing before downstream staining. All gels will expand overnight in DI water before imaging.

Detailed quantification of the expansion factor and quality control procedures are provided in the Supplementary Information.

### Dual-mode multiscale imaging platform

A dual-mode imaging platform integrating OPM and ASLM was developed to enable coordinated multiscale volumetric imaging of expanded liver tissue. The imaging system includes two complementary light-sheet systems. The ASLM path provides ∼460 nm isotropic resolution over a field of view of 774 µm x 435 µm using remote focusing and axial sweeping, ideal for high-detail subcellular imaging^18^. The OPM module delivers a 3 µm lateral resolution over an effective 5.3 mm x 3.8 mm FOV, suitable for rapid, low-resolution screening and target identification^28^. Both modules share a common sample holder designed to support expanded gels up to 20x their original size.

The ExOPM imaging module took inspiration from DvOPM^28^. In this design, the illumination path consists of a laser source with 4 wavelengths (405nm, 488nm, 561nm, 642nm) combined into a single fiber output, coupled into a Protected Silver Reflective Collimator (Thorlabs RC08FC). To ensure even illumination across the lateral dimensions of the light sheet, the light passed through a Powell Lens with a 10° angle (LaserLine Optic) to collimate it in 1 dimension by an achromat doublet, L1 (f=60mm, Thorlabs), and focused on a light sheet in the other dimension. The illumination beam was directed onto a 4kHz resonant galvometric mirror (Novanta), enabling light-sheet pivoting to reduce shadow artifacts due to absorption in the specimen^29^. The light sheet is then focused on the sample using a simple relay lens pair, L2 and L3, with a slit to control the numerical aperture (NA) of the illumination (f= 100mm, f=125mm respectively, Thorlabs.) With a 6mm input beam diameter at the 125mm focusing lens, it creates an effective illumination NA of 0.024.The detection assembly consisted of a Mitakon Speedmaster 65 mm f/1.4 photographic lens acting as the primary objective (O1), followed by a filter wheel, a 90° folding mirror, a Nikon AF-S NIKKOR 85 mm f/1.4G lens serving as the secondary objective (O2), and a CMOS camera, the Ximea MU196MR, for image acquisition The photographic lens pair were chosen such that the ratio of its focal length (85mm/65mm) create a magnification, M = 1.31, that closely match the refractive index of water (n=1.33) to form an aberration-free image in the remote space. Although the lens combination is capable of supporting a theoretical detection numerical aperture (NA) of approximately 0.36, the effective system NA was limited to 0.20 by the 32 mm clear aperture of the filter wheel.

A comprehensive description of the ASLM imaging setup and acquisition parameters can be found in our previous publication^18^. Additional information on the imaging system is provided in Supplementary Information, and the image acquisition parameters are listed in Supplementary Table 2.

Regions of interest (ROIs) for high-resolution ASLM imaging were selected based on low-magnification OPM volumetric images. The central vein and portal vein were manually identified from the OPM overview to define liver zonation^30,31^, and the microscope was then switched to ASLM to image cells within the corresponding zonal regions. This workflow ensured that all cells included in the analysis originated from the designated liver zones. No ROIs were excluded after acquisition unless image quality was insufficient for reliable segmentation and quantitative analysis. All images used for analysis were raw images and were not subjected to deconvolution.

### Deconvolution

Images corresponding to the zoomed-in regions in Fig. 1E–G were processed using a custom deconvolution pipeline implemented in MATLAB. The method combines blind with Lucy-Richardson deconvolution. A synthetic point spread function (PSF) was first generated based on the imaging parameters, such as excitation/emission numerical aperture, excitation/emission wavelengths, and voxel sampling size. We then used blind deconvolution (deconvblind) in MATLAB with ten iterations to estimate the data PSF from the acquired image data. Next, the retrieved PSF from the blind deconvolution was used for image restoration with the Lucy-Richardson deconvolution routine (deconvlucy) in MATLAB, also using ten iterations. Final deconvolved image stacks together with the retrieved PSFs were exported as volumetric TIFF files for subsequent analysis. We limited the number of iterations to ten to avoid clipping of dim features and over-deconvolution. To reduce boundary artifacts during deconvolution (both blind and Lucy-Richardson deconvolutions), raw image volumes were symmetrically padded in all three spatial dimensions. After deconvolution, image volumes were cropped back to their original dimensions, intensity-normalized to match the native dynamic range of the raw data and converted to 16-bit format for downstream visualization and quantitative analysis. The MATLAB version is not critical for our deconvolution pipeline; versions from 2019 to 2025 were tested and yielded the same results.

### Single-cell segmentation and global morphology analysis

Individual hepatocytes were segmented from 4x expanded volumetric ASLM datasets using a multiview segmentation pipeline integrating orthogonal-view 2D segmented stacks with pretrained micro-sam^32^ models and 3D label fusion using u-Segment3D^33^, as illustrated in Supplementary Fig. S2. Prior to segmentation, image volumes underwent background correction, percentile-based intensity normalization, slice-wise contrast enhancement using CLAHE (clip limit = 0.01), and Gaussian smoothing (σ = 1.0 for whole-cell segmentation). Cells were segmented using the micro-SAM ViT-B LM model from three orthogonal views (XY, XZ, and YZ) followed by u-Segment3D fusion to generate 3D segmentations. Small, disconnected objects (<50 voxels for cells) were removed. 3D surface meshes were extracted from segmented cell masks for downstream quantitative analysis using u-Unwrap3D^34^.

Global morphological features, as shown in Supplemental Table 1, including volume, surface area, sphericity, solidity, compactness^35,36^, aspect ratio, and surface roughness, were extracted from each cell using MATLAB R2023b (MathWorks) based on our previously described geometric analysis^37,38^. What is noteworthy is that inter-featured correlation analysis revealed some morphological parameters were highly correlated (Fig. S3B). Therefore, only relatively uncorrelated features were selected for downstream analysis in Fig. 2G. Principal component analysis (PCA) was performed on shared morphological features across Normal, CTNNB1, and NRAS groups to identify dominant axes of structural variation. Before PCA, all features were standardized by z-score normalization to ensure equal weighting across variables with different scales. To further assess potential batch or biological sample-dependent variability, PCA was additionally performed with sample identity annotation, confirming that the observed variation was not primarily driven by sample-specific effects (Fig. S3A, additional analyses are provided in the Supplementary Information). PCA was subsequently computed using singular value decomposition, and the first two principal components (PC1 and PC2) were used for downstream visualization and analysis. Feature contributions to the PCA space were quantified from PC1 and PC2 loading magnitudes to identify the dominant morphological descriptors underlying group differences. PC1 scores for individual cells were compared to Normal, CTNNB1, and NRAS groups using one-way ANOVA followed by Tukey’s HSD multiple-comparison test. For visualization, PC1 distributions were displayed as boxplots with overlaid single-cell data points.

### Membrane curvature analysis

The membrane mean curvature was quantified from 3D single-cell surface masks using u-Unwrap3D^34^. For each segmented cell, a triangular surface mesh, *S* (*x, y, z)*, was extracted from the binary cell mask using marching cubes with Gaussian pre-smoothing (presmooth =1.0) and CGAL-based remeshing to improve mesh quality. Mean curvature was computed using a gradient smoothing parameter of 5.0 to reduce sensitivity to mesh noise. For each cell, vertex-wise curvature values were exported and summarized using distribution-based metrics, including the 95th percentile of absolute mean curvature, to capture high-curvature membrane features such as protrusions, irregular edges, and local surface roughness. These per-cell curvature measurements were then used for downstream comparison of membrane morphology across normal, CTNNB1-activated, and NRAS-activated hepatocytes.

### Mitochondrial segmentation and morphology analysis

Mitochondria were segmented from volumetric ASLM datasets using a multistep 3D segmentation pipeline. Briefly, prior to segmentation, image volumes underwent background correction, percentile-based intensity normalization, slice-wise contrast enhancement using CLAHE (clip limit = 0.01), and Gaussian smoothing (σ = 0.9 for mitochondrial segmentation). Then individual image slices were first segmented using the micro-SAM^32^ human-in-the-loop framework to generate high-accuracy 2D mitochondrial masks. The slice-wise segmentation information from orthogonal views were subsequently integrated using the indirect fusion method implemented in u-Segment3D to reconstruct volumetric 3D mitochondrial masks (Fig. S5F). Each mitochondrion was represented as an individually labeled object within the final 3D image stack. Before quantitative analysis, a preprocessing and cleanup step was performed to remove small, disconnected fragments and segmentation artifacts (small objects <20 voxels will be removed), After that, the mitochondrial segmentation accuracy was evaluated by comparing reconstructed 3D mitochondrial masks across XY, XZ, and YZ orientations. Segmentation masks were resized to raw image dimensions using nearest-neighbor interpolation. Accuracy was quantified on a slice-by-slice basis using Intersection-over-Union (IoU) and Dice similarity coefficient metrics between reconstructed masks, reference masks, and foreground regions estimated from raw fluorescence images using Otsu thresholding. In addition, segmentation boundaries were overlaid onto raw fluorescence images for qualitative visual validation of mask accuracy and structural continuity throughout the 3D volume (Fig. S5D-E).

Morphological measurements were then computed independently for each mitochondrial object using voxel-calibrated 3D analysis. Extracted features included mitochondrial volume, principal axis lengths, mean diameter, elongation ratio, and additional shape descriptors derived from 3D region property analysis implemented in scikit-image *regionprops*. To further characterize mitochondrial network organization, skeleton-based analyses were performed to quantify branching architecture and tubular connectivity. Each mitochondrial object was skeletonized in three dimensions using the *skimage.morphology.skeletonize_3d* algorithm. Skeleton refinement was subsequently applied to remove short branches (terminal branches shorter than 1 μm and side branches shorter than 2 μm) likely introduced by segmentation noise or local reconstruction artifacts. Following refinement, the longest endpoint-to-endpoint geodesic path within each mitochondrial network was identified and retained as the major mitochondrial backbone for downstream structural analysis.

Additional morphology descriptors were derived by integrating volumetric and skeletal measurements. Cross-sectional area was estimated as:

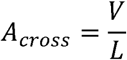

where represents mitochondrial volume and represents total skeleton length. Average mitochondrial diameter was estimated assuming cylindrical geometry:

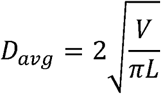

These measurements enabled quantitative assessment of mitochondrial size, tubular elongation, branching complexity, and network organization across different oncogenic conditions. For quality control, filtered mitochondrial label volumes and corresponding backbone skeleton reconstructions were exported as 3D TIFF files for visual inspection. A representative example is shown in Fig. S6G. In addition, because total mitochondrial skeleton length exhibited substantial variability and measurement noise across individual mitochondria (Fig. S6E, F), the mitochondrial population was stratified into three length-based fractions. The longest fraction (top one-third by skeleton length) was subsequently selected for downstream visualization and analysis, as highly elongated mitochondria more reliably capture biologically relevant differences in mitochondrial network organization and structural remodeling. Besides, feature correlation analysis of the mitochondrial measurements revealed redundancy among multiple morphological parameters (Fig. S6B). Therefore, mitochondrial diameter and volume were selected as the primary quantitative features, as they capture complementary aspects of mitochondrial ultrastructure and provide robust descriptors of mitochondrial morphological remodeling.

### Multiscale feature integration and classification

Multiscale structural integration was performed using quantitative features extracted from global cellular morphology, membrane curvature, and mitochondrial ultrastructure across Normal, CTNNB1, and NRAS hepatocyte populations. Global morphology features included sphericity, surface area, solidity, aspect ratio, roughness, volume, compactness, and long-axis length, whereas membrane curvature was represented by the 95th percentile and median mean curvature. Mitochondrial architecture was quantified using average mitochondrial diameter, mitochondrial volume, and total mitochondrial skeleton length.

To integrate measurements acquired from independent imaging datasets, pseudo-bulk samples were generated separately for each genotype. Each pseudo-bulk consisted of randomly sampled measurements from global cell morphology (4 cells), membrane curvature (4 cells), and mitochondrial morphology (120 mitochondria). Sampling was performed with replacement to enable repeated generation of pseudo-bulk samples from the finite pool of observations while preserving the underlying feature distributions within each genotype. Within each modality, the mean and standard deviation of each feature were calculated to generate aggregated descriptors of cell morphology, membrane topology, and mitochondrial organization (Fig. S6H). These modality-specific descriptors were then concatenated into a single multiscale feature vector representing the structural state of each pseudo-bulk. For classification analysis, 40 pseudo-bulk samples were generated per genotype for both the training and testing datasets in each cross-validation fold. For PCA visualization, 50 pseudo-bulk samples were generated per genotype. Random sampling was performed using NumPy’s random number generator (*default_rng*), with the random seed defined as *repeat x 100 + fold* index to generate independent pseudo-bulk samples for each cross-validation iteration.

Prior to dimensionality reduction and classification, selected size-related features (Volume, Compactness, Surface Area, Long Length, Average Mitochondrial Diameter, Mitochondrial Volume, and Total Skeleton Length) were log-transformed prior to z-score normalization. To balance the contributions of different structural scales, modality-specific weighting factors were applied (global morphology = 1.0, membrane curvature = 1.7, mitochondrial morphology = 1.7). For integrated models, the top 14 features were selected using an ANOVA F-test before classifier training. Principal component analysis (PCA) was then performed on the integrated multiscale feature vectors to visualize structural separation between oncogenic states. For visualization only, pseudo-bulk samples corresponding to the most extreme 10% of distances from each genotype centroid were excluded to improve display clarity, whereas all pseudo-bulk samples were retained for classification analyses.

Supervised classification was performed using Random Forest classifiers trained on either individual modality feature sets or the integrated multiscale feature set. Random Forest models were constructed with 400 decision trees, unlimited tree depth, a minimum leaf size of one, and all remaining parameters set to their default values. Classifier performance was evaluated using repeated three-fold GroupKFold cross-validation. Group assignments were defined by the original biological sample identifier (*SampleID*), ensuring that all measurements originating from the same sample remained within a single fold. GroupKFold partitioning was performed before pseudo-bulk construction, and training and testing pseudo-bulks were generated independently from their respective data pools. Consequently, no original cells, curvature measurements, or mitochondria were shared between training and testing datasets, thereby preventing data leakage. Classification performance was quantified using the macro-averaged F1 score across 15 independent cross-validation repeats, and feature importance was estimated from the trained Random Forest models and averaged across all cross-validation iterations. Statistical comparisons among classification strategies were performed using one-way ANOVA followed by Tukey’s multiple-comparison test.

### Data visualization and statistical analysis

All plots were generated using matplotlib and seaborn. PCA plots, violin plots, boxplots, scatter plots, heatmaps, and classifier performance summaries were generated using consistent group color schemes across all figures: Normal (gray), CTNNB1 (pink), and NRAS (blue).

Uniform Manifold Approximation and Projection (UMAP) visualization was performed using the Python umap-learn implementation based on five mitochondrial morphological features (mitochondrial volume, minor axis length, average diameter from volume, elongation ratio, and major axis length). Prior to dimensionality reduction, features were standardized by z-score normalization using StandardScaler. Two-dimensional embeddings were generated with n_neighbors = 50, min_dist = 0.05, spread = 1.0, n_components = 2, the default Euclidean distance metric and random_state = 42. Group-level UMAP positions were represented by the median UMAP1 and UMAP2 coordinates, and pairwise group separation in the embedding was quantified as the Euclidean distance between group median coordinates. For comparison, Euclidean distances were also calculated between group median vectors in the original standardized five-dimensional feature space.

For multigroup comparisons, statistical significance was evaluated using one-way analysis of variance (ANOVA) followed by Tukey’s honestly significant difference (HSD) post hoc multiple-comparison test. For comparisons involving two independent groups, including mitochondrial morphology quantification, statistical significance was assessed using a two-sided Mann–Whitney U test. Statistical significance was defined as P < 0.05 (*), P < 0.01 (**), P < 0.001 (***), and P < 0.0001 (****). Data are presented as mean ± s.d. unless otherwise indicated.

## Acknowledgements

K.M.D. acknowledges support from the Cancer Prevention and Research Institute of Texas (RP250571), the National Cancer Institute (U54CA268072), and the National Institute of General Medical Sciences (RM1GM145399). R.F. receives support from the National Cancer Institute (U54CA268072), the National Institute of General Medical Sciences (R35GM133522), and the National Institute of Biomedical Imaging and Bioengineering (R01EB035538). H.Z. is supported by the National Institute of Diabetes and Digestive and Kidney Diseases (DP1DK139976) and the National Institute on Alcohol Abuse and Alcoholism (R01AA028791). H.M-F. is supported by the National Cancer Institute (K99CA283246). OpenAI’s ChatGPT was used to edit author-generated text for clarity, organization, and flow; all scientific content and final wording were reviewed and approved by the authors.

## Author Contributions

Q.S., H.Z., and K.M.D. conceived and designed the study. Q.S. led day-to-day execution of the study, coordinated experimental and analytical contributions across the team, developed and optimized experimental workflows, performed imaging experiments, led data analysis, and prepared the original manuscript draft. T.N. and H.M.-F. contributed to methodology development and writing of the original draft. K.W. contributed methodology, resources, animal tissue preparation, tissue isolation, and writing of the original draft. K.B. performed cloning and contributed to methodology development. F.Y.Z. and Z.S. performed segmentation-based image analysis. B.-J.C. and R.F. assisted in image deconvolution. X.W. contributed image acquisition software development. J.H. contributed to data curation and data presentation. H.M.B. optimized sample labeling methods. H.Z. provided resources, visualization, and supervision.

K.M.D. provided resources, visualization, supervision, project administration, and funding acquisition. All authors reviewed and edited the manuscript.

## Data and code availability

Image analysis code used in this study is available in a public GitHub repository at https://github.com/NicoleShenv587/Multiscale-Morphology-Analysis-of-Oncogenic-Alterations. The microscope control software, navigate, is publicly available at https://github.com/TheDeanLab/navigate. Detailed design files and assembly documentation for the OPM system are available at https://github.com/TheDeanLab/altair. Archived versions of all software will be deposited in Zenodo upon publication. Owing to the size of the volumetric imaging datasets, a representative subset of the data will be archived upon publication. The remaining imaging data are available from the corresponding author upon request.

