## Supplemental Materials for "Deciphering the Multiscale Morphology of Somatic Oncogenic Alterations in Hepatocellular Carcinoma"

**Extended methods:**

**Acquisition of Large-Scale ExOPM Image Volumes**

To demonstrate fast large-scale volumetric imaging, an expanded liver specimen was imaged over a volume measuring 3.75 mm x 9.8 mm x 400 µm (post-4x expansion). The data was acquired using ExOPM in 1.5h using 10 µm step size across 2 channels with α-tubulin and mitochondria staining, enabling visualization of the overall distribution of activated hepatocytes across the tissue while preserving up to subcellular structural information.

**Calibration of the spatial offset between ExOPM and ExASLM modules**

The displacement between the ExOPM and ExASLM imaging systems was manually determined using sparsely distributed 4x expanded spheroids as fiducial calibration targets. After the focal planes of both imaging modules had been independently aligned and fixed, an isolated spheroid was centered within the ExOPM field of view using the shared motorized X-, Y-, and Z-stage, and the corresponding stage coordinates were recorded. The imaging mode was then switched to ExASLM, and the same spheroid was manually translated until it was centered within the ExASLM field of view. The new stage coordinates were recorded, and the coordinate differences between the two imaging positions were calculated to determine the fixed translational offset between the two microscope configurations. The calibration was repeated for ten additional isolated spheroids, and the average offset was used to establish the coordinate transformation for accurate automated navigation and multiscale image registration between the ExOPM and ExASLM imaging modes.

**Expansion factor quantification and quality test**

To evaluate expansion isotropy, expansion factors were quantified at both the bulk gel level and the subcellular level. Gel expansion was first assessed by measuring dimensional changes before and after expansion. Pre-expansion tissues (1x) and first-round expansion (4x) gels were imaged using a smartphone under top illumination, whereas second-round expansion (16x) gels were imaged using a Bio-Rad ChemiDoc MP Imaging System (Fig. S1B). To assess expansion across the full tissue thickness, side-view volumetric images were acquired using a custom-built Oblique Plane Microscopy[34] and subsequently stitched in ImageJ. Gel dimensions were compared along all three orthogonal axes to evaluate volumetric isotropy (Fig. S1D). To further validate isotropic expansion at the subcellular scale, nuclear dimensions were measured at each expansion stage. Nuclear sizes were quantified in both XY and XZ views and compared across expansion conditions to determine the consistency of subcellular expansion in lateral and axial dimensions (Fig. S1F). Measured expansion factors showed low variability across samples, consistent with the nominal expansion factor. Therefore, all measurements were normalized using the nominal expansion factors (4x and 16x), which introduced negligible errors relative to the observed biological differences.

#### Permutation Test for Assessing Biological Label Structure

To quantitatively assess separation significance, multinomial logistic regression models were trained using five-fold stratified cross-validation (Fig. S3C-D). Classification accuracy obtained using the true genotype labels was compared against a null distribution generated through permutation testing. Specifically, cell-type labels were randomly shuffled while preserving the original feature matrix, and for each permutation, the complete cross-validation and classification procedure was repeated to generate a null distribution of classification accuracies. The observed accuracy obtained using the true labels was then compared against the null distribution to calculate a one-sided permutation p-value according to:

$$p = \frac{1 + \sum_{i=1}^{N} I(a_i \geq a_{obs})}{N + 1}$$

where $a_{obs}$represents the observed classification accuracy, $a_i$denotes the accuracy obtained from the $i$-th permutation, and $N$is the total number of permutations performed. This approach verified that classification performance depended on biologically meaningful label structure rather than random feature separation.

#### Correlation and feature association analysis

Pairwise relationships among morphological and mitochondrial features were assessed using Pearson correlation coefficients computed from standardized measurements (Fig. S3B and S6B). Correlation matrices were visualized as heatmaps to identify coordinated structural remodeling associated with oncogenic activation. For selected feature comparisons, linear regression analysis and coefficient of determination ($R^2$) values were additionally calculated.

### Supplementary Figures

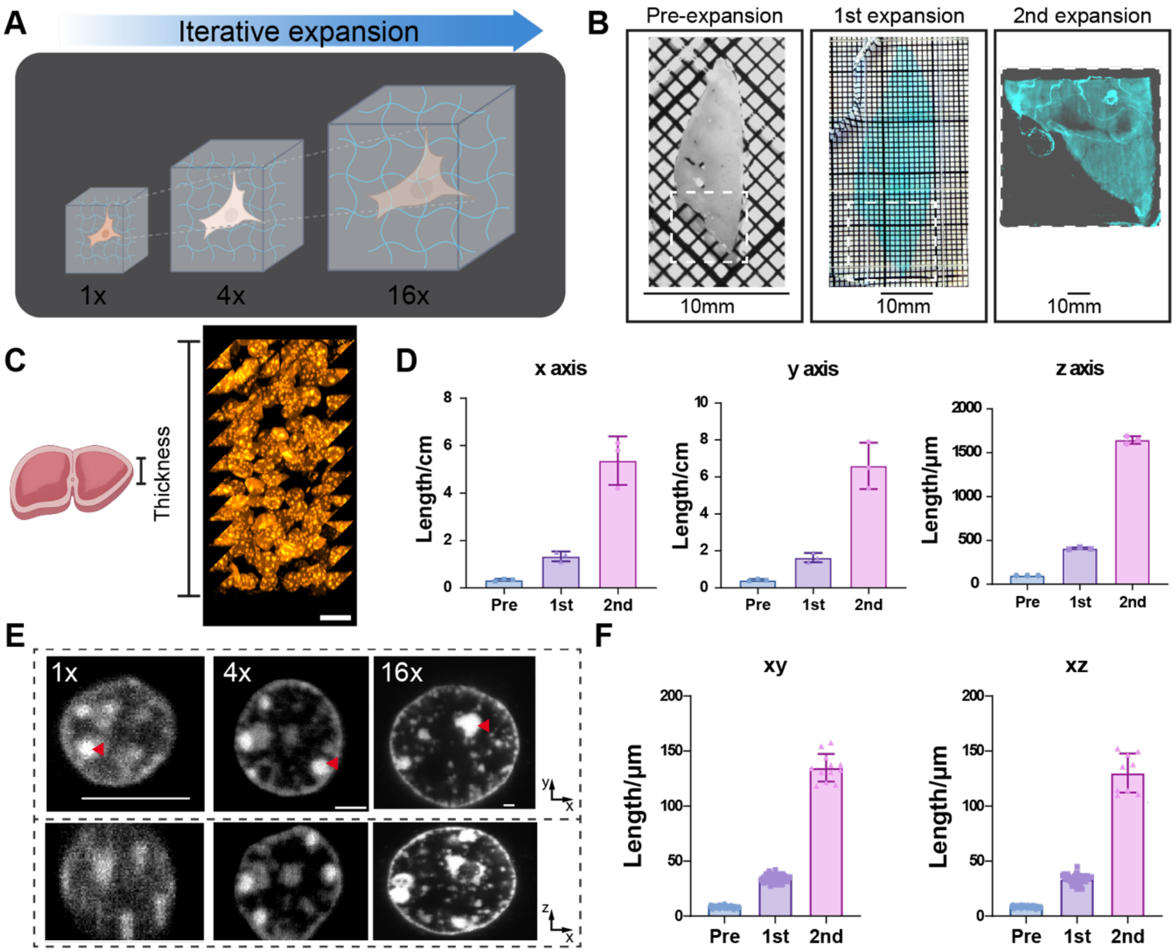


**Figure S1. Iterative expansion microscopy enables isotropic 3D scaling across tissue and subcellular levels.** (A) Schematic of the iterative expansion workflow, illustrating three rounds of gel embedding to achieve sequential 4x ($1^{st}$) and 16x ($2^{nd}$) expansion. (B) Representative gel images at each stage of expansion: pre-expansion (1x), $1^{st}$ expansion (4x) and $2^{nd}$ expansion(16x). Gels are stained with NHS-647 after expansion. (C) Full-thickness view of a sample after the $1^{st}$ expansion (4x). Scale bar, 50 µm (post-expansion) equivalents to 12.5 µm (pre-expansion). (D) Measurements of sample dimensions along the x, y, and z axes across expansion stages. (E) Nuclear IF imaging (SYTOX staining) showing resolution at each expansion stage using ASLM. Red arrows indicate nucleoli. Scale bar, 10 µm. (F) Quantification of nuclear diameter in xy and xz dimensions for each expansion stage.

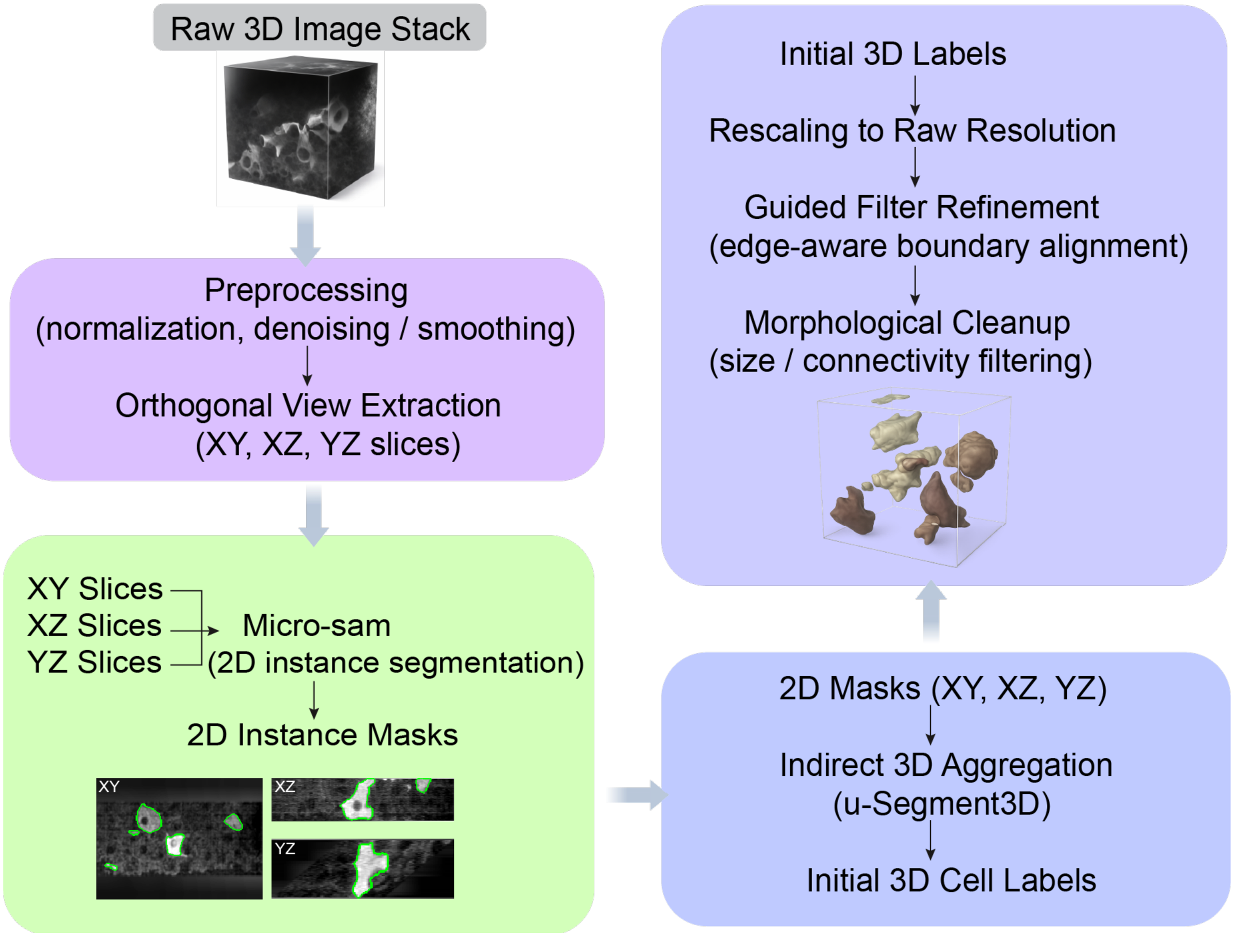


**Figure S2. Schematic of the single-cell 3D segmentation workflow.** Raw volumetric images are preprocessed and resampled to generate orthogonal views (XY, XZ, YZ), followed by 2D instance segmentation and fusion into 3D cell masks. Postprocessing includes label refinement, artifact removal, and boundary alignment.

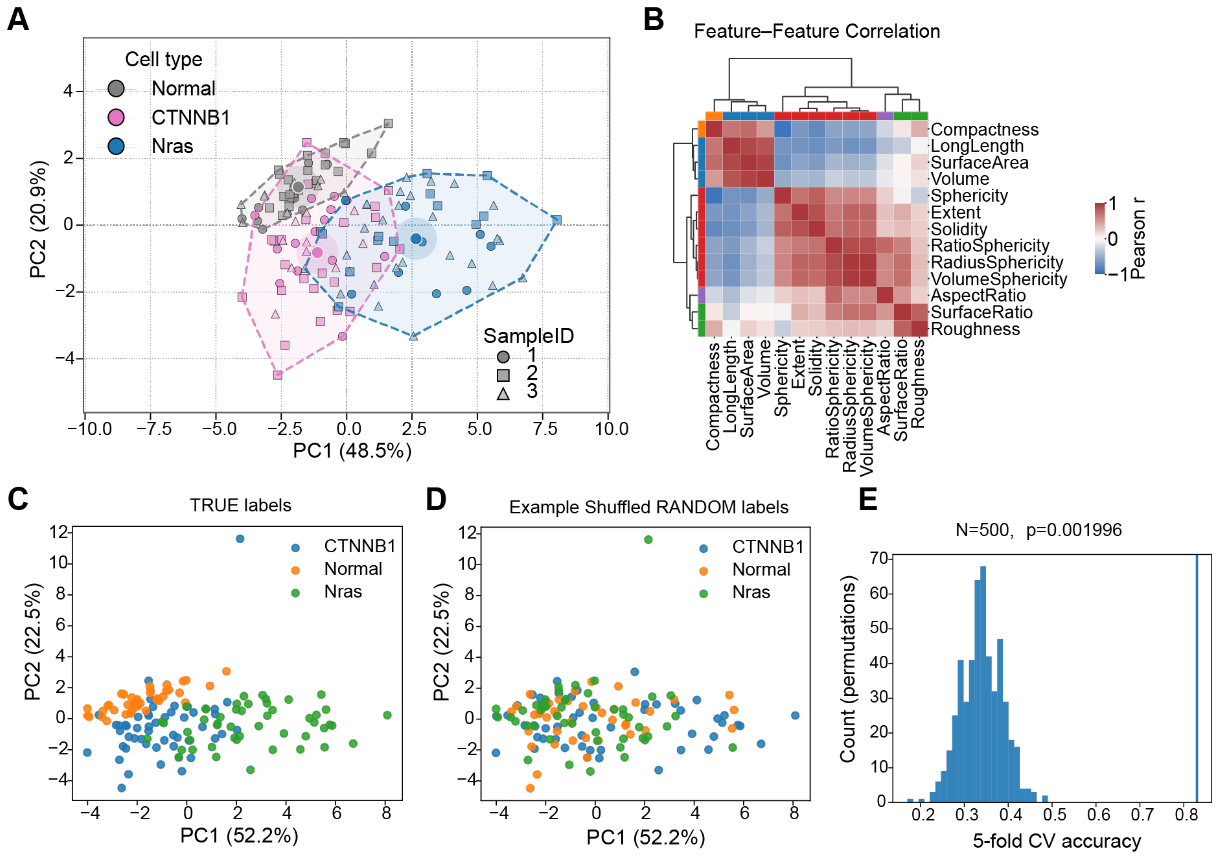


**Figure S3. Validation and characterization of single-cell morphological features after oncogenic activation**. (A) Principal component analysis (PCA) of single-cell morphological features, with points annotated by sample identity. (B) Feature–feature correlation matrix for global morphology metrics. (C) PCA of cellular morphological features colored by true labels (Normal, CTNNB1, NRAS). (D) PCA of the same dataset with randomly permuted labels. (E) Permutation test of classification accuracy using multinomial logistic regression with 5-fold cross-validation. Histogram shows the distribution of accuracies from shuffled labels, with the observed accuracy from true labels indicated by a vertical line. The p-value indicates the probability of achieving equal or higher accuracy under random label assignment.

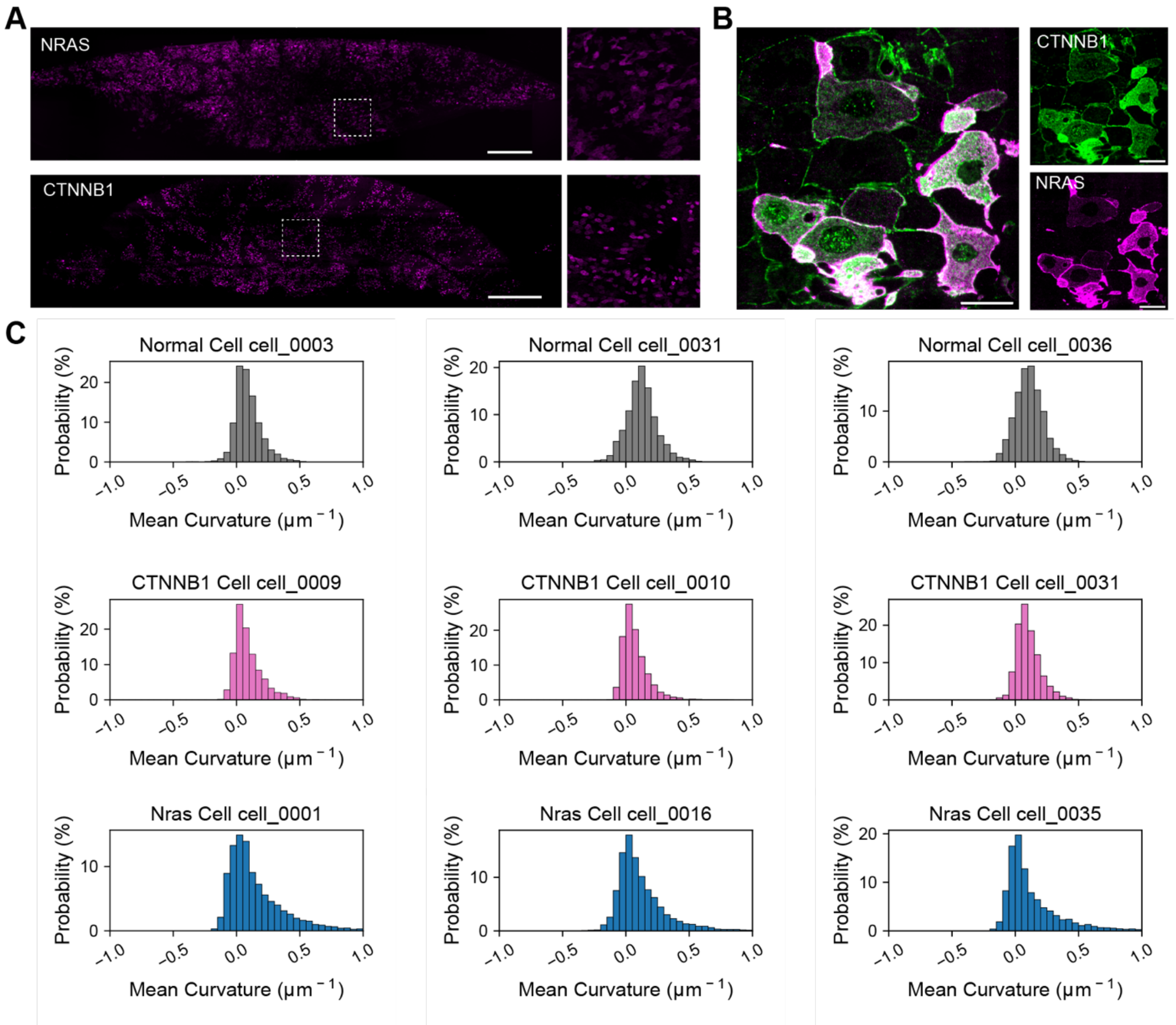


**Figure S4. Spatial distribution and curvature analysis of oncogene activated hepatocytes.** (A) Whole tissue view showing the distribution of NRAS- and CTNNB1-activated cells (magenta) within the liver. Right panels show zoom in regions from the boxed areas in left panels. Scale bars, 5,000 μm (post-expansion) and 1,250 μm (pre-expansion). (B) Immunofluorescence image of a double-mutant sample, with CTNNB1 labeled in green and NRAS in magenta. Scale bars, 50 μm (post-expansion) and 12.5 μm (pre-expansion). (C) Representative distributions of cell membrane curvature for Normal (gray), CTNNB1 (pink), and NRAS (blue) hepatocytes.

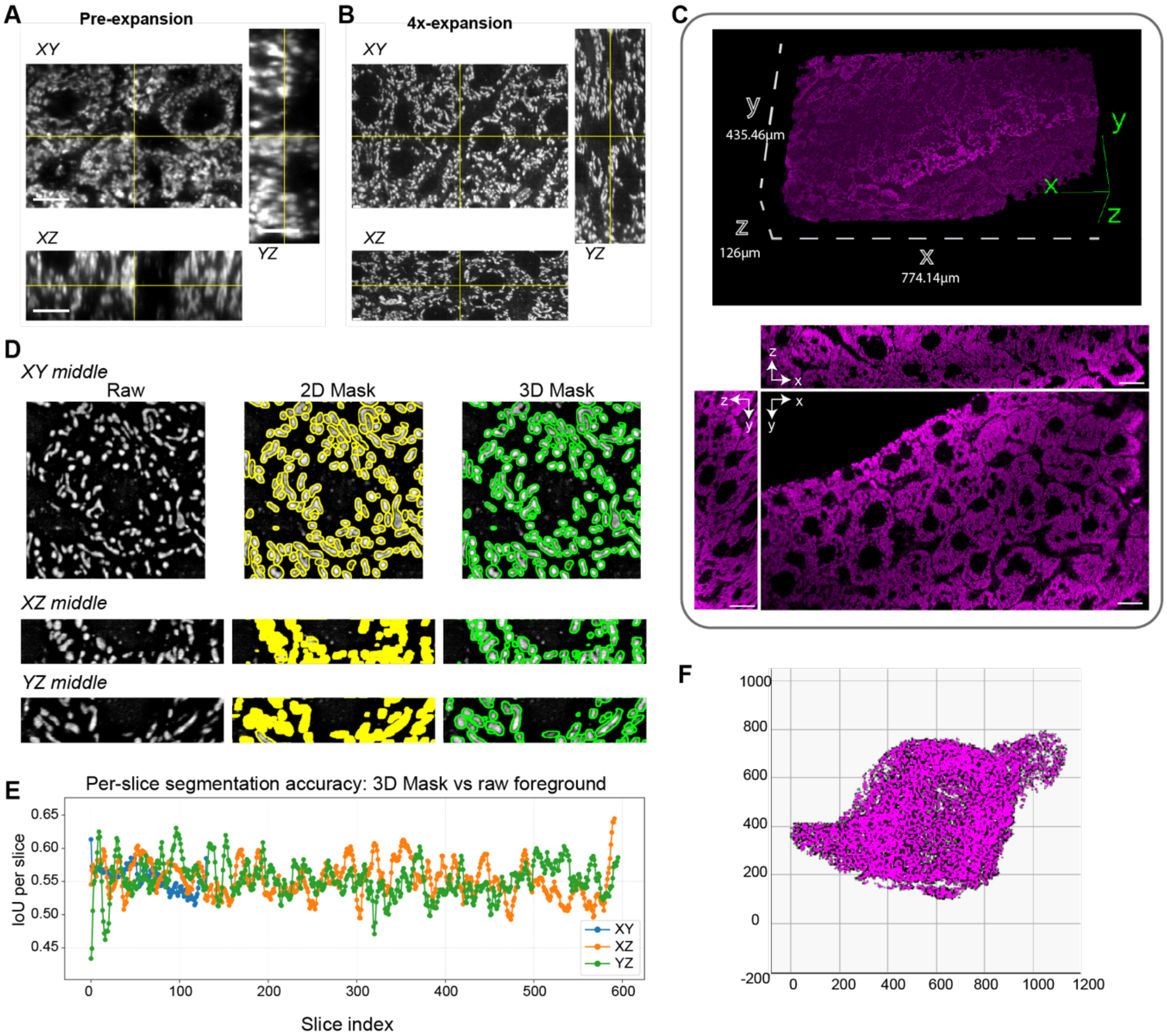


**Figure S5. 3D mitochondrial imaging and validation of segmentation accuracy in expanded liver tissue.** (A, B) Orthogonal 3D views of mitochondria in liver tissue before expansion (A) and after 4x expansion (B), acquired by ASLM. Mitochondria are labeled with Hsp60. Scale bars, 10 µm. (C) 3D rendering and orthogonal views of 4x expanded liver tissue showing mitochondrial organization. Scale bars, 50 µm (post-expansion) and 12.5 µm (pre-expansion). (D) Evaluation of 3D segmentation accuracy across orthogonal views; left panels show representative middle slices of the raw image (grey) in XY, XZ, and YZ orientations, and middle and right panels show overlays of segmentation boundaries from 2D segmentation (yellow) and reconstructed 3D segmentation (green). (E) Slice-wise intersection-over-union (IoU) between reconstructed 3D masks and raw image foreground, estimated by intensity-based thresholding. Lines represent individual orientations (XY, XZ, YZ). (F) Example of aggregated 3D mitochondrial segmentation within a single cell.

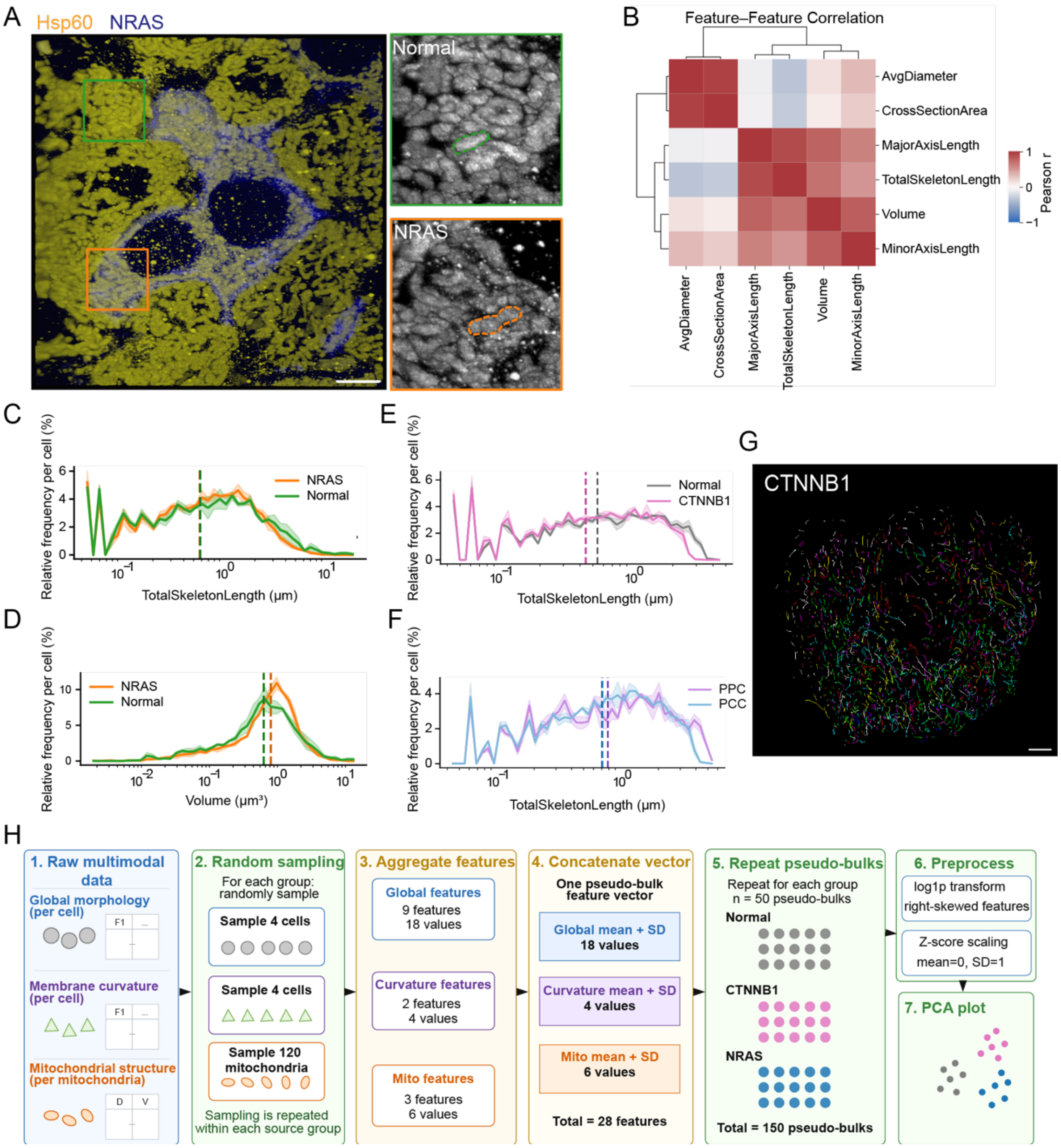


**Figure S6. Mitochondrial organization and quantitative feature analysis across oncogenic states.** (A) Immunofluorescence (IF) image of 4x expanded NRAS sample showing NRAS expression (blue) and mitochondria labeled with Hsp60 (yellow). Zooms in regions highlight mitochondrial organization in NRAS⁺ (orange) and NRAS⁻ (green) cells. Desh lines indicate the boundary of a single mitochondria in each condition. (B) Feature–feature correlation matrix of mitochondrial measurements. (C, E, F) Distributions of total mitochondrial skeleton length normalized per cell for Normal versus NRAS (C), Normal versus CTNNB1 (E), and periportal (PP) versus pericentral (PC) regions (F). (D) Distribution of mitochondrial volume normalized per cell for NRAS and Normal cells. (G) Computed mitochondrial skeletons for individual mitochondria within a representative CTNNB1-mutant cell. Scale bars, 10 μm (post-expansion) and 2.5 μm (pre-expansion). (H) Schematic workflow for pseudo-bulk generation by random aggregation of cell morphology, curvature, and mitochondrial features within each group (n = 50 per group).

### Supplementary Tables

| Feature | Definition | Expression | Interpretation |
|---|---|---|---|
| Volume | Total voxel volume of the object | V | Overall object size |
| SurfaceArea | Surface area of the triangulated boundary | A | Boundary complexity & size |
| Sphericity | Similarity to a perfect sphere (1 = sphere) | π^(1/3) (6V)^(2/3) / A | Higher = more spherical |
| Solidity | Fraction of convex hull volume occupied | V / V_convex | Lower = more concave |
| LongLength | Maximum convex hull vertex distance | max \|\|p_i - p_ j\|\| | Global elongation |
| Aspect Ratio | Shortest to longest distance ratio | min(L1,L2,L3) / max(L1,L2,L3) | 1 = isotropic |
| Roughness | Surface irregularity vs convex hull | A / A_convex | >1 indicates corrugation |
| Extent | Volume occupancy of bounding box | V / V_bbox | Compactness in bbox |
| SurfaceRatio | Surface area vs bounding sphere | A / (4πR_c²) | Normalized surface |
| VolumeSphericity | Volume vs bounding sphere | V / (4/3 πR_c³) | Sphere filling |
| RadiusSphericity | Equivalent radius vs bounding radius | (EquivDiameter/2) / R_c | Size-normalized |
| RatioSphericity | Inscribed-to-circumscribed ratio | R_i / R_c | Roundness |
| *Compactness | Surface-to-volume compactness | A^(3/2) / V | Higher = less compact |

**Symbols**

$V$ = object volume
$R_{eq}$ = equivalent sphere radius
$A$ = object surface area
$L_1, L_2, L_3$ = axis lengths
$R_c$ = circumscribed sphere radius
$R_i$ = inscribed sphere radius

*Compactness of a 3D object in computer vision is often defined using inverse surface-volume ratios to amplify deviations from ideal compact shapes and improve sensitivity to irregular geometry.

**Table 1. 13 features included in the global cell morphology analysis pipeline.**

| | ExOPM | ExASLM |
|---|---|---|
| Laser Power | • 488nm - 100mW<br>• 642nm - 100mW | • 488nm - 40 mW<br>• 561nm - 100 mW<br>• 642nm - 100 mW |
| Exposure (ms) | **50** | **80** |
| Z-step size (µm ) | **10** | **0.2** |
| Voxel size ($\mu m^3$ ) | 1.06 x 1.06 x 10 | 0.17 x 0.17 x 0.2 |
| Illumination objective lens NA | 0.02 | 0.7 |
| Detection objective lens NA | 0.18 | 0.7 |
| Emission Filter | • 488nm - FF01-515/30-32<br>• 561nm - FF01-595/31-32<br>• 642nm - FF01-670/30-32 | • 488nm - FF01-515/30-32<br>• 561nm - FF01-595/31-32<br>• 642nm - FF01-670/30-32 |

**Table 2. Image Acquisition Parameters for ExOPM and ExASLM Imaging**

| Structural analysis | Group | Mice | ROIs | Cells | Mitochondria |
|---|---|---|---|---|---|
| Global cell morphology | Normal | 3 | 10 | 42 | – |
| | CTNNB1 | 3 | 29 | 50 | – |
| | NRAS | 3 | 40 | 49 | – |
| Membrane curvature | Normal | 3 | – | 20 | – |
| | CTNNB1 | 3 | – | 18 | – |
| | NRAS | 3 | – | 22 | – |
| Mitochondrial morphology | Normal | 3 | – | 3 | 6,715 |
| | CTNNB1 | 3 | – | 3 | 6,269 |
| | NRAS | 3 | – | 3 | 7,195 |
| Zonation mitochondrial morphology | Periportal (PP) | 3 | – | 3 | 2,197 |
| | Pericentral (PC) | 3 | – | 3 | 4,895 |

*Membrane curvature was quantified using the same segmented cell population analyzed for global morphology. Because curvature estimation is highly sensitive to surface mesh quality, cells with incomplete, fragmented, or artifact-containing surface meshes that failed quality control were excluded prior to curvature analysis.

**Table 3. Summary of biological replicates and quantified objects used for multiscale structural analyses.**
